# Rampant convergent evolution of vascular oddities and a synnovation characterize the rapid radiation of Paullinieae lianas

**DOI:** 10.1101/2025.07.12.664521

**Authors:** Israel L. Cunha Neto, Zachary Kozma, Isaac H. Lichter-Marck, Pedro Acevedo-Rodríguez, Joyce G. Onyenedum

**Author notes:** **Author Contributions:** J.G.O., I.L.C.N., Z.K., P.A.R. designed research; J.G.O. and P.A.R. provided samples; J.G.O., I.L.C.N., Z.K., P.A.R. performed research; J.G.O., I.L.C.N., Z.K., I.H.L-M. contributed analytic tools; J.G.O., I.L.C.N., Z.K., P.A.R. analyzed data; J.G.O., I.L.C.N., Z.K. wrote the paper, with additional contributions by P.A.R. and I.H.L-M. **Competing Interest Statement:** The authors declare no competing interest.

## Abstract

Climbing plants have independently evolved thousands of times and are particularly successful in tropical forests, yet their anatomical and evolutionary distinctiveness remains poorly understood. Among the most striking innovations in woody climbers—or “lianas”—are vascular variants: modifications to xylem and phloem that depart from the typical growth found in trees and shrubs. In this work, we leverage the fourth largest lineage of neotropical lianas, Paullinieae (Sapindaceae), to elucidate the evolution of development of vascular variants, and to test key innovation hypotheses in this megadiverse group. We reconstruct the largest phylogeny of any liana lineage to date (227 species, 351 nuclear genes), revealing a rapid radiation in the Miocene. Our anatomical evaluation of 462 species uncovered six patterns of vascular variants spanning three developmental categories—procambial, cambial, and ectopic cambia—which evolved repeatedly across the tree. Using stochastic mapping and a developmental complexity framework, we show that evolutionary transitions from typical growth disproportionately favored developmentally simple variants, suggesting that developmental accessibility constrains macroevolutionary trajectories. Despite the temporal overlap between the disparification of vascular variants, and the diversification rate shift, we find no evidence that vascular variants alone drive species diversification. Instead, diversification rates correlate with the presence of tendrils, climbing growth forms, and zygomorphic flowers. These results suggest a synnovation— a suite of synergistic innovations rather than a single trait—as the driver of lineage radiation in Paullinieae. Our study highlights how integrating phylogenomics, developmental anatomy, and trait evolution can illuminate the evolutionary mechanisms shaping plant diversity.

**Significance Statement:** Lianas are functionally distinct from trees and are rising in abundance across neotropical forests. Yet, it remains poorly understood how these plants are developmentally constructed, and what traits underlie their success. Here, we present the largest molecular phylogeny of any climbing plant lineage and show that vascular variants—unusual stem anatomies that enhance flexibility— are remarkably diverse and have evolved repeatedly in Paullinieae. However, these striking modifications are not linked to species diversification. Instead, we find that the combination of three traits—the climbing habit, coiling tendrils, and zygomorphic flowers—collectively drove the diversification of Paullinieae. Our findings challenge traditional views of key innovations, showing that synergistic trait combinations, rather than any single trait, can catalyze evolutionary radiations and shape biodiversity.

## Introduction

One of the most conspicuous, yet understudied, evolutionary innovations is the emergence of climbing plants. Unlike self–supporting trees and shrubs, climbing plants rely on other plants for physical support as they ascend through the forest canopy in search of light (1). Despite the complexity of this phenotype, climbing plants have independently evolved thousands of times across the plant tree of life, where the climbing habit has been hypothesized to be a key innovation driving species diversification in flowering plants (2, 3). In complement to their evolutionary success, the past twenty years of ecological research has converged on an alarming finding that demands further investigation: many woody climbers––i.e., “lianas”––are responding to increasing forest fragmentation, drought, and rising temperatures, by growing larger and more abundant, particularly in the neotropics (4, 5). Therefore, the increasing dominance of lianas poses a threat not only to the trees upon which they grow but also to forest–level carbon sequestration capacities (5). With these new insights, the question emerges: what makes lianas so distinct (6), given they are made from the same raw materials (cells and tissue types) as self-supporting trees and shrubs?

Within the stems of many lianas, we find unique anatomical modifications that render these woody stems adapted to climb (6). These developmental tweaks manifest as reconfigurations of the vascular tissues (xylem and phloem) that deviate from the “typical” growth common to nearly all seed plants (7). These deviations are collectively termed “vascular variants” (8). Vascular variants enhance the mechanical resistance, hydraulic properties, and flexibility of lianas (6, 9). Thus, vascular variants are likely a key ingredient in the ability of lianas to climb.

The diversity of vascular variants—both in form and developmental origins—is highest within the monophyletic tribe Paullinieae (Sapindaceae) (10, 11), the fourth largest lineage of neotropical lianas (Fig. 1A-E), for which previous studies have recognized 10 distinct patterns (Fig. 1F–N) observed in five of the seven genera (12). As pointed out in earlier studies focusing on individual genera (13–15), this diversity likely indicates a case of repeated evolution in Paullinieae at large. Within Sapindaceae, a single evolutionary transition from self-supporting growth forms to lianas characterized Paullinieae, which diversified to encompass 25% of the family’s species. However, the drivers of this remarkable diversification process remain unknown (16). A likely contender is the evolution of vascular variants, given that this trait is exclusive to Paullinieae within Sapindaceae (16). However, Paullinieae also exhibits other potential key innovations that have been hypothesized to drive speciation, including coiling tendrils, zygomorphic flowers (in all genera except *Thinouia*), and the climbing habit itself (Fig. 1B–E) (16). Taken together, Paullinieae is an ideal system to elucidate both the fundamental processes shaping the evolutionary lability of vascular development and the underlying drivers of species diversification in climbing plants. Yet, the lack of a robust phylogeny and comprehensive morpho-anatomical survey has limited our ability to fully realize this potential.

**Figure 1.**
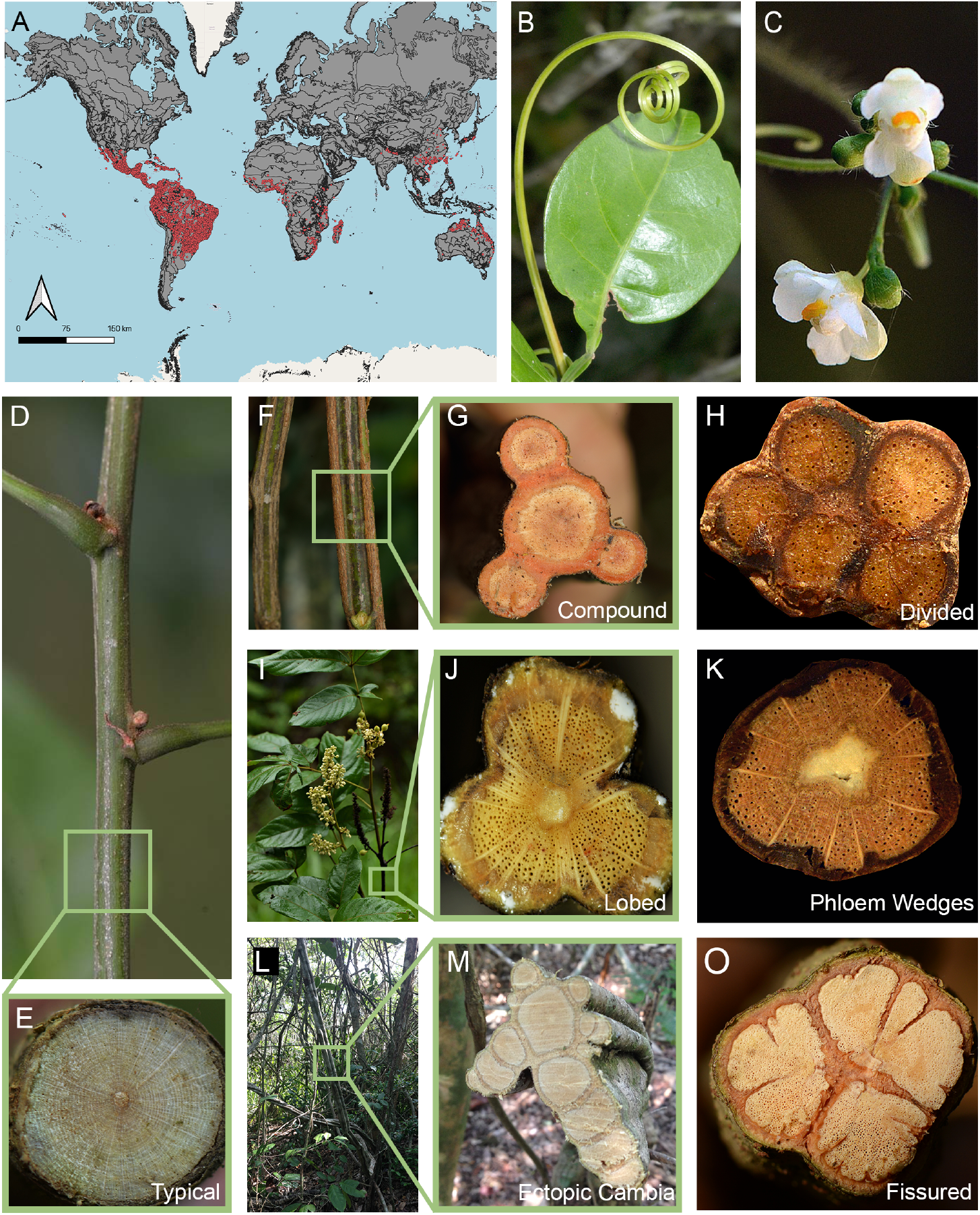
Geographical distribution and morphological traits characterizing Paullinieae lianas. (*A)* Distribution occurrence points acquired from BIEN (Botanical Information and Ecology Network) in maroon. *(B)* Tendril and leaflet of a trifoliolate leaf; *Serjania paucidentata. (C)* Zygomorphic flower; *Cardiospermum corindum.* (*D-E*) Stem with typical growth, formed by a cylindrical body of wood and bark derived from a single vascular cambium; *P. bilobulata.* (*F–O*) Stems with vascular variants, including procambial variants (*F–H*), cambial variants (*I–M, O*), and ectopic cambia (*L-M*). Species name: (*F-G*) *P. pinnata. (H) S. erythrocaulis*. (*I-J*) *P. stellata.* (*K*) *S. mexicana*. (*L-M*) *Thinouia restingae*. (*N*) *Urvillea laevis*. Image credit: B-K: Pedro Acevedo-Rodríguez (61); *L-M*: Israel L. Cunha Neto; *N:* reproduced from Cunha Neto et al. (14), with permission from Oxford University Press. Stem diameters range from 20 to 50 mm.

In this study, we leverage the Paullinieae tribe to elucidate the evolution of development of vascular variants and to test the drivers of diversification in this important neotropical lineage. First, we developed an essential, but lacking resource: the largest time calibrated phylogeny of a liana lineage, to date. With this chronogram, we ask two key questions: (1) What is the tempo and mode of evolution of development of vascular variants in this megadiverse group? And (2) what morphological traits are driving species diversification? By integrating systematics, anatomy, and phylogenomics, our study provides insights into both developmental and evolutionary mechanisms underlying the ecological and evolutionary success of lianas.

## Results and Discussion

### Nuclear DNA Dataset Generates a Robust Phylogeny

In this study, we reconstructed a time-calibrated tree of Paullinieae, yielding the most comprehensive phylogeny for the tribe. This phylogenomic framework provides the necessary evolutionary context to understand the macroevolution of this megadiverse group. Our final dataset included 333 samples and 227 species, encompassing 212 Paullinieae and 15 tree/shrub outgroups from supertribe Paulliniodae and members of other Sapindaceae lineages, including *Cupania* spp., *Blomia prisca, Guindilia trinervis,* and *Talisia* spp. (*SI Appendix,* Table S1–S2). We retrieved a range of 127 to 352 genes targeted by the Angiosperms353 probe set (17), and the final dataset, obtained from the coding sequences (CDS), comprised 351 genes (*SI Appendix,* Fig. S1; see Table S3 for assembly statistics). This dataset was used to infer the phylogeny using a partitioned Maximum Likelihood (ML) analysis of the concatenated supermatrix with IQ–TREE (18) (*SI Appendix*, Fig. S2), and the 351 individual gene trees were utilized to construct a species tree with ASTRAL (19) (*SI Appendix*, Fig. S3). The IQ–TREE and ASTRAL results both recovered robust phylogenies, for which all but one node for major clades is well–supported (BS = 100, PP = 1) (Fig. 2A; *SI Appendix*, Figs. S2, S3). A comparison of these two trees reveals that the primary differences are within infra-generic relationships, with no changes in the backbone topology (*SI Appendix*, Fig.S4).

**Figure 2.**
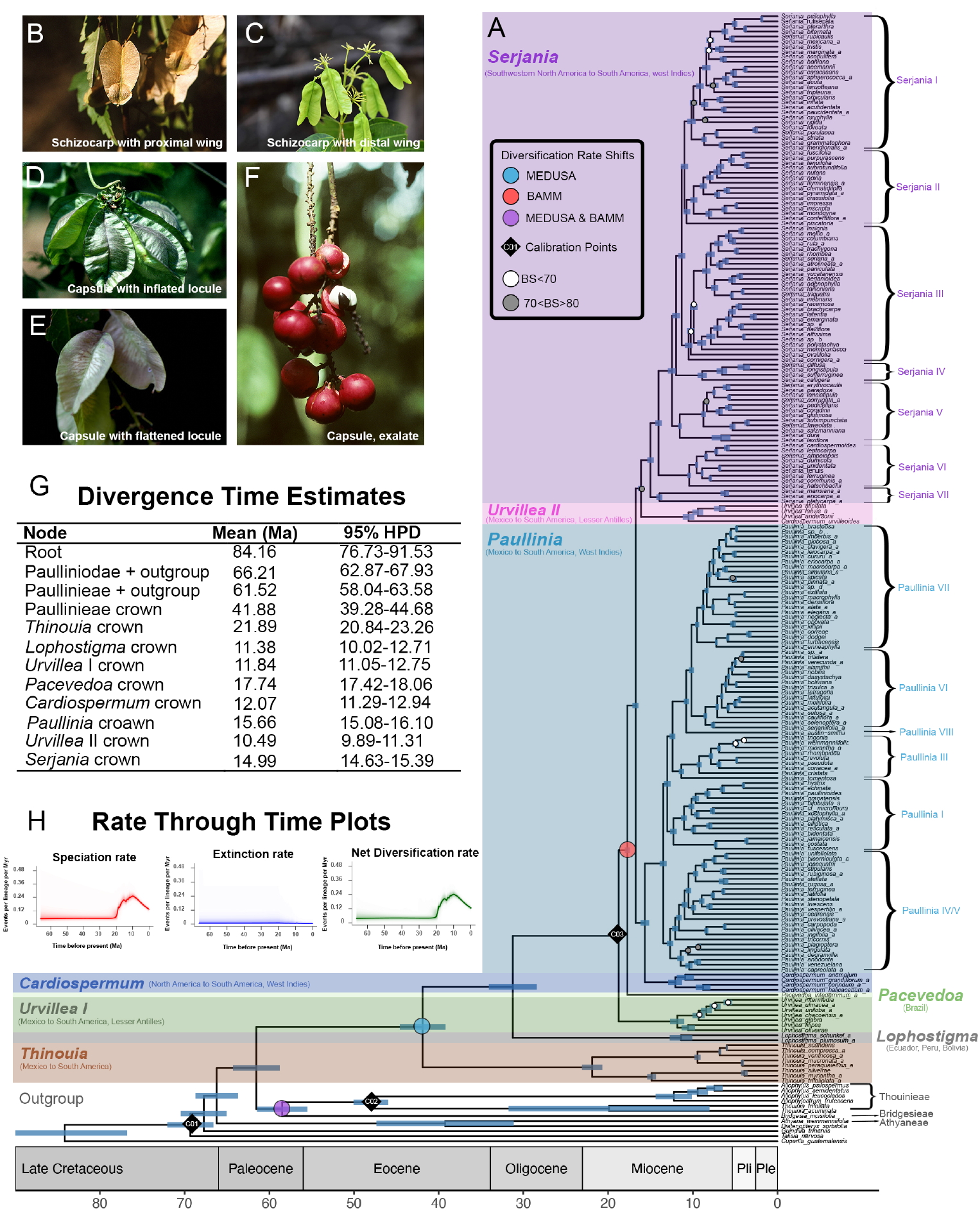
Time-calibrated phylogeny of Paullinieae generated with TreePL displaying eight major clades, node ages, and fruit diversity. (*A)* Chronogram including one sample per species presented with main nodes indicating bootstrap values (all 100 unless stated otherwise). The infrageneric clades of *Paullinia* and *Serjania* are indicated. *Paullinia* follows a clade numbering system based on Chery et al. (21), with modifications reflecting updated phylogenetic relationships: Formerly separate clades IV and V now form a single monophyletic group, clade II is absent in our updated phylogeny, *P. austin smithii* is assigned to a newly defined clade, clade VIII, expanding Chery et al. (21) original seven clades. (*B–F*) Fruit types. (*B)* Schizocarpic with proximal wing; *Lophostigma plumosum.* (*C*) Schizocarpic with distal wing; *Thinouia myriantha.* (*D)* Capsule with inflated locule; *Urvillea chacoensis.* (*E)* Capsule with flattened locule; *U. laevis.* (*F*) Capsule, exalate; *Paullinia spicata*. (*G*) Summary of divergence time estimates for the major clades; the tree is time-calibrated with two secondary calibrations and one fossil. (*H*). Rate-through-time plots for Paullinieae from BAMM analysis. Image credit: *B–F*: Pedro Acevedo-Rodríguez (61).

Our newly constructed phylogenies include eight major clades that correspond to the six traditionally recognized genera of Paullinieae, except for the split of *Urvillea* into two clades (*Urvillea* I and *Urvillea* II), and the newly described genus *Pacevedoa* (20), dismembered from the polyphyletic *Cardiospermum* (Fig. 2). *Thinouia* is strongly supported as sister to the rest of Paullinieae (Fig. 2A). After the divergence of *Thinouia*, we found a successively nested topology as follows: *Lophostigma, Urvillea I*, then *Pacevedoa.* Subsequently, we found two sister clades, one formed by *Urvillea* II and *Serjania* with relatively low support (BS = 77) and the other including *Paullinia* and *Cardiospermum* (Fig. 2A). Within each major clade, most clades are well supported (BS > 80), with a few exceptions in *Urvillea* I, *Paullinia* and *Serjania* which have moderate (70< BS <80) or low support (BS < 70) (Fig. 2A).

The confirmation of monophyly for four genera (*Paullinia, Serjania, Lophostigma*, and *Thinouia* (14, 16, 20–23)) validated that genus-level morphological synapomorphies, as defined by Radlkofer (11) still hold true today. However, the polyphyly of two genera (*Cardiospermum* and *Urvillea*) revealed that the usually go-to trait “fruit type” (capsule vs. schizocarp) is unreliable in delimiting the genera. Nevertheless, our two *Urvillea* clades are broadly separated by nuanced fruit traits: *Urvillea* I has fruits with inflated locules and three seeds per fruit (Fig. 2B–F; (24)), while *Urvillea* II mostly has fruits with flattened locules. Our phylogeny identified three instances of paraphyletic species, which are recognized as either part of a known species complex (*P. weinmannifolia* and *P. alata* and their allies) or are involved in taxonomic questions (*T. myriantha*) related to newly delimited species (i.e., *T. silveirae* (23)). The phylogeny presented in this work represents a substantial advancement to Paullinieae phylogenetics, whereby previous molecular studies either had moderate species sampling and a low quantity of molecular marks (14, 16, 20– 22), or had low species sampling and a high quantity of molecular markers (25, 26). Thus, our newly constructed tree provides a foundation for additional taxonomic revisions and the elucidation of micromorphological and/or genomic features to further delimit the boundaries of *Cardiospermum* and *Urvillea* (*sensu lato*).

### The Burst in Species Diversity in Paullinieae is Correlated with Shifts in Diversification Rates

To understand the tempo and mode of diversification in Paullinieae, we time-calibrated our maximum likelihood tree with one primary and two secondary calibrations (Fig. 2A; *SI Appendix,* Fig. S5). Our chronogram demonstrated a “broom-and-handle” tree shape, characteristic of a recent and rapid radiation. Paullinieae lianas diverged from their tree/shrub relatives ∼66 Ma, then experienced a 40 Myr lag before a diversification rate shift in the Miocene ∼18 Ma, culminating in > 50% of extant species (Fig. 2G).

To determine whether the Paullinieae radiation is linked to changes in diversification rates, we performed diversification analyses using two methods. Both MEDUSA and BAMM detected a net diversification rate shift in one outgroup genus, *Allophylus*, comprised of ∼200 trees and shrubs (Fig. 2A). Within the Paullinieae lianas, MEDUSA detected a diversification rate shift at the most recent common ancestor of *Pacevedoa* and the rest of Paullinieae (Fig. 2A; *Appendix*, Fig. S6) and BAMM detected one in the ancestor of *Urvillea* I and the rest of Paullinieae (Fig. 2A; *SI Appendix*, Fig. S7). Our diversification analyses and rate–through–time plots indicate that the Paullinieae radiation was the result of increasing speciation rates coupled with constantly low extinction rates (Fig. 2H). The recent radiation of Paullinieae corroborates the finding that during the Eocene-Oligoene climatic shift, the putative ancestors of Paullinieae migrated from Southeast Asia and Australia to South America through Antarctica, colonizing the warmer regions of the New World, where species numbers proliferated in the Miocene (27). Interestingly, the Paullinieae radiation is contemporaneous with several groups of angiosperms undergoing similar radiations (e.g., Crassulaceae, Lecythidaceae (28, 29), including lineages containing lianas, such as Annonaceae and Bignoniaceae (30, 31). Our findings indicate an earlier divergence than reported by Urdampilleta et al. (20), who found the supertribe Paulliniodae originates in the early Eocene (∼51 Ma), while Paullinieae originated in the late Eocene (∼37 Ma), respectively. Since similar calibration points were utilized across both studies, our discrepancies are likely explained by differing taxonomic sampling (in this study, 333 samples vs. 91) and more molecular markers (in this study, 351 vs. 3). Additionally, the discrepancy in dates may arise from varying calibration methods: TreePL in our study versus BEAST in the previous phylogeny.

To better understand the potential drivers of rate shifts in Paullinieae, we began by characterizing, then elucidating the evolution of one of the most distinct attributes of the tribe: the diversity of vascular variants.

### Transition Rates to Different Vascular Variants Vary by Developmental Complexity

To characterize the diversity of vascular biology in Paullinieae, we surveyed 462 of the 463 species by evaluating the stem anatomy from herbarium specimens, image databases, liquid-preserved specimens, and the literature (*SI Appendix*, Table S4). We identified that Paullinieae species are nearly equally split between those with typical growth (n=232) or a vascular variant (n=230) (*SI Appendix*, Table S4). While the variants may appear superficially random, the six observed patterns can be binned into three categories (Fig. 3; *SI Appendix*, Fig. S8, S9), based on the timing of developmental modifications *sensu* Cunha Neto (8). Within the procambial variant category (i.e., alterations during primary growth), we find two patterns: the compound (151 species) and divided (8 species). Within the cambial variant category (i.e., alterations during secondary growth), we find three patterns: the lobed (30 species), phloem wedges (26 species), and fissured (2 species). The ectopic cambia category houses the ectopic cambia pattern, characterized by the formation of additional cambia in unusual locations (13 species) (Fig. 3A–I; Fig. S8, S9). This level of diversity is striking even among lineages with vascular variants, which usually have one or two patterns (e.g., Bignoniaceae, Malvaceae; (32, 33)), or an impressive diversity of patterns, albeit restricted to one or two categories (e.g., Apocynaceae, Fabaceae, Malpighiaceae) (8).

**Figure 3.**
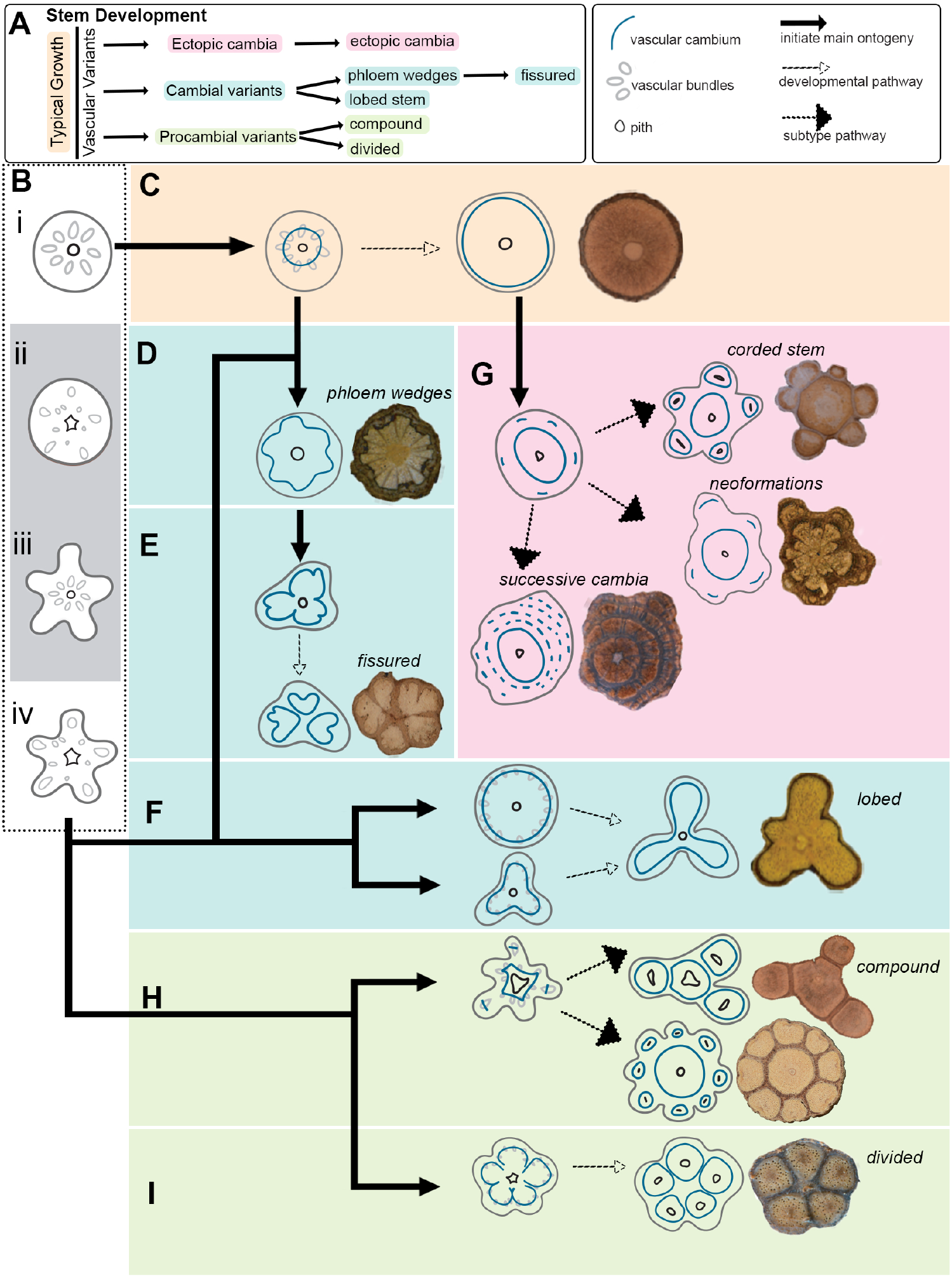
Diversity of stem development and vascular variants in Paullinieae lianas. (*A*) Hierarchical developmental pathways leading to typical growth and vascular variants. (*B*) The initial developmental stage for the different ontogenies maybe one of the following: (i) circular shape and typical eustele (single ring of vascular bundles), (ii) circular shape and atypical eustele (irregularly distributed bundles), (iii) lobed shape and typical eustele, or (iv) lobed shape and atypical eustele. (i) and (iv) are more common, and (ii) and (iii) are less common. (*C)* Typical development begins with a circular shape and a typical eustele (i), from which a single vascular cambium arises to produce secondary xylem (wood) and secondary phloem (inner bark). (*D–I*) Ontogenies for the six patterns of vascular variants; Zstem development may start from typical growth or atypical growth (as described in *B*); (*D*) Phloem wedges: sectors of the cambium produce higher amounts of secondary phloem in comparison to the secondary xylem; this process creates phloem wedges, which can vary in depth and width. (*E*) Fissured: the vascular system is dissected due to continuous production of vascular tissues in some sectors of the cambium, and reduced activity in the sectors where phloem wedges are formed. (*F*) Lobed: The vascular cambium produces both secondary xylem and secondary phloem at higher rates in some sectors, and reduced rates in other sectors; this process creates lobes and furrows, which can vary in number and shape. (*G*) Ectopic cambia: arise is stems after some secondary growth, in which a new cambium arises repeated times and create additional vascular tissues; new cambia can form complete cylinders (top), embedded vascular units in the cortex or vascular system (middle), or continuous arcs/rings (bottom); given their similar development, all three subtypes can be grouped under the term ectopic cambia. The image in the middle also represents a case of a combination of phloem wedges + ectopic cambia. (*H*) Compound: Multiple cambia arise from the irregular distribution of vascular bundles in primary growth; each set of dislocated vascular bundles forms an independent cylinder in addition to the central cylinder; the peripheral cylinders can originate from individual dislocated bundles (top, Compound I) or multiple dislocated bundles (bottom; Compound II). (*I*) Divided: vascular bundles are irregularly distributed, which leads to independent vascular cambia generating vascular cylinders. Species name: *(C) Paullinia echinata; (D) Serjania reticulata*; *(E) Urvillea laevis; (F) P. caloptera*; *(G) Thinouia scandens* (top), *S. grandifolia* (center); *S. pernambucensis* (bottom). *(H) S. caracasana* (top), *P. pinnata* (bottom); *(i) S. paleata.* Image credit: *C–F, G (top* and *center*), *H–I*: Pedro Acevedo-Rodríguez (Acevedo et al., 2015, onwards); G (bottom): reproduced from Cunha Neto et al. (62), with permission from The Linnean Society of London. Drawings not to scale; stem diameters range from 10 to 30 mm.

To elucidate the evolutionary history of Paullinieae’s unique diversity, we employed stochastic character mapping. To our surprise, we found that the ancestral state of Paullinieae, and all eight major clades, was reconstructed as having typical growth (*SI Appendix*, Fig. S11, S12). Thus, Paullinieae experienced rampant convergent evolution of compound, phloem wedges, lobed, fissured, and ectopic cambia––all patterns that are found in more than one genus (*SI Appendix*, Fig. S11, S12). Moreover, we found additional complexity within Paullinieae, including subtypes of compound and ectopic cambia (Fig. 3), species with more than one pattern (*SI Appendix*, Fig. S10), and confirmed the existence of polymorphic species (*SI Appendix,* Fig. S11; Marques et al., 2025)).

To further probe the nature of the convergent evolution of vascular variants, we studied their ontogenies, identifying that the three variant categories fall along a spectrum of developmental complexity: (1) procambial variants are difficult to construct, requiring the reorganizing of the number, location and/or polarity of vascular bundles to build the primary vascular system; (2) cambial variants are intermediate, requiring only modifications to the relative abundance of xylem versus phloem of a single cambium, and (3) ectopic cambia, which produce multiple cambia in the same stem, may be the easiest to construct, as it is the most common variant across plants (8, 34), partially explained by simple genetic tweaks inducing the downregulation of *KNOX* genes (35). With this developmental framework, we leveraged our dataset to test whether developmentally easy variants have higher transition rates, and vice versa.

Across 1000 stochastic character maps, we found that the most common evolutionary transitions from typical growth were to the cambial variants, phloem wedges (7.71), and lobed (5.35). The second most common transitions were to the ectopic cambia (5.25). Finally, the least common transitions were procambial variants: compound (2.67) and divided (0.88). The only exception was the infrequent transitions to fissured (0.9) (Fig. 4B). When investigating reversals from each pattern back to typical growth, we did not find a clear pattern; thus, the loss of variants appears to be independent of developmental construction and seemingly stochastic. On the other hand, the evolutionary transitions from typical growth to diverse patterns mirror the spectrum of developmental complexity, with higher transitions toward easier developmental patterns.

**Figure 4.**
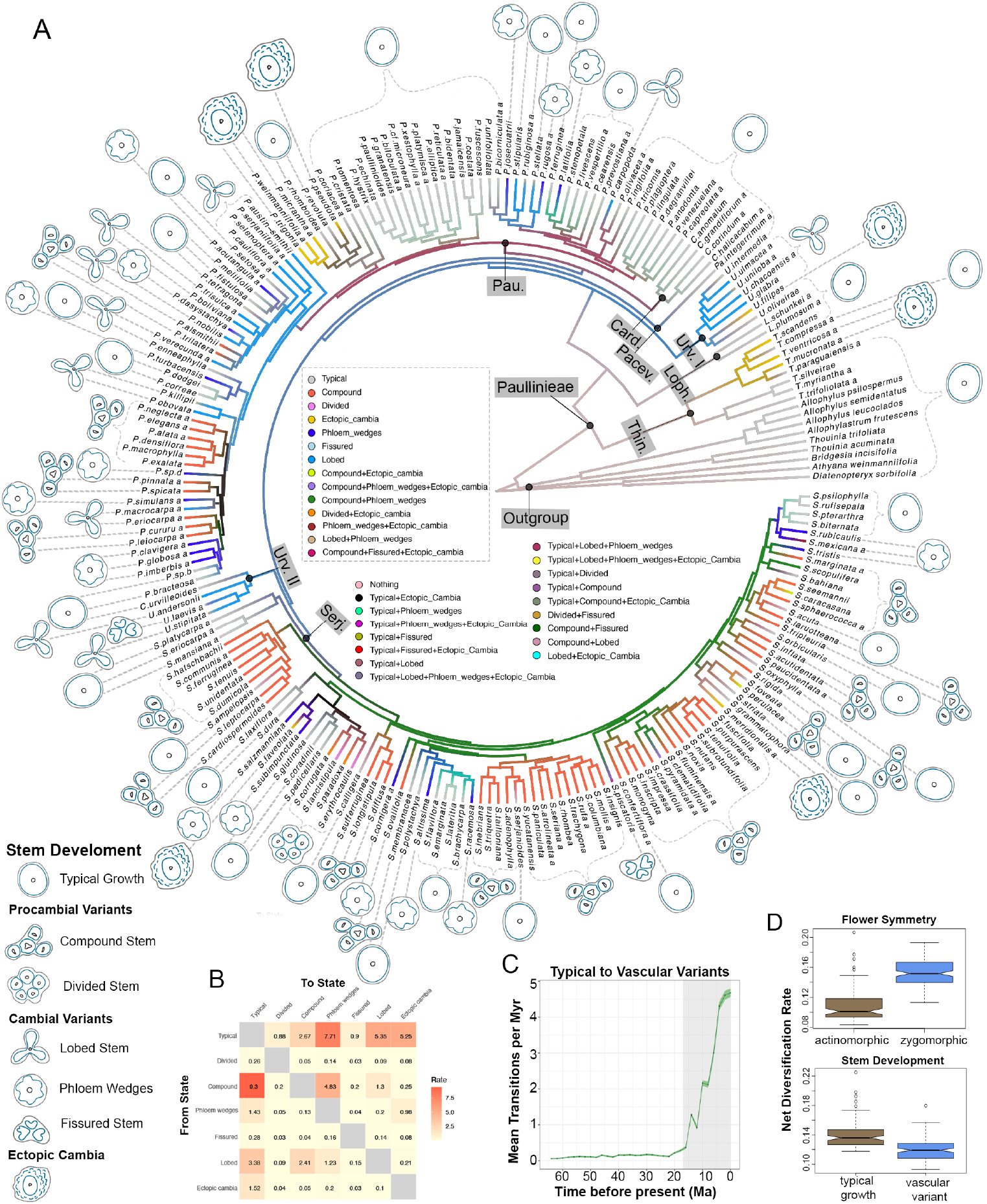
Stochastic character mapping showing inferred evolutionary history of vascular variants using and diversification rate shifts. (*A*) Overlap of 1000 stochastic character maps under All Rates Different (ARD) transition model PARAMO illustrating all possible developmental combinations.. (*B–C*) Timing and number of transitions from typical growth to vascular variants, extracted from 1000 simulated stochastic histories using the patterns analysis (*SI Appendix,* Fig. S13). *(B)* Heatmap of evolutionary transitions of stem development vascular variants. (*C*) Line plot showing the transitions from typical to vascular variants through evolutionary time. *(D)* Model averaging from hidden state speciation and extinction (HiSSE) models shows significantly different diversification rates for flower symmetry and the presence or absence of vascular variants.

To understand when this rampant convergent evolution of vascular variants occurred, we extracted geological time and estimated transitions from our stochastic mapping (*SI Appendix*, Figs. S12). This plot revealed an exponential increase in evolutionary transitions from typical growth to vascular variant patterns during the Miocene (Fig. 4C), which corresponds to the recent radiation of the tribe. This finding led us to ask whether the rapid disparification of vascular variants, or other morpho-anatomical traits, correlates with the shift in diversification in Paullinieae.

### The Acquisition of Lianas, Tendrils, and Zygomorphic flowers––Not Vascular Variants–– Influence Diversification Dynamics

Given our finding that Paullinieae experienced a recent and rapid radiation, coupled with the fact that these lianas possess a suite of traits hypothetically linked to diversification, we had the unique opportunity to test key innovation hypotheses. We began by testing whether the evolution of vascular variants may have driven species diversification in Paullinieae. While the increase in species diversification and the disparification of vascular variants temporally overlapped in the Miocene, our HiSSE analysis (*SI Appendix,* Tables S5-S6) showed no evidence of a link between transitions to vascular variants and the diversification heterogeneity in this lineage (best fit model= CID4; *SI Appendix,* Tables S7). This result is not entirely surprising, as both large (e.g., *Paullinia*) and small (e.g., *Urvillea sensu lato*) genera in Paullinieae include many species with vascular variants (13, 14). Moreover, vascular variants are a vegetative trait; thus, they are not directly linked to creating and/or reinforcing reproductive isolation. Cunha Neto et al. (36, 37) also found that vascular variants are not strongly responsible for diversification processes in Caryophyllales. In these studies, a procambial variant (medullary bundles) was correlated with diversification shifts, with a preference for the BiSSE model over the others in one of the families within this large order (36). However, when the correlation between medullary bundles and species diversification was performed at the order level, that hypothesis was not confirmed (37).

If not vascular variants, then what traits underlie the rapid radiation of Paullinieae? The literature posits that the climbing habit (2), tendrils (38), and zygomorphic flowers (27) are all candidate key innovations––these traits are all present in Paullinieae. To test these hypotheses, we applied the HiSSE model, which uncovered that each of these traits partially explains the diversification heterogeneity in Paullinieae (*SI Appendix,* Table S6). The results of our model– fitting analysis indicated that the HiSSE model provided the best fit for the climbing mechanism and growth forms, indicating that transitions to lianas and tendrils are implicated in the diversification (*SI Appendix,* Tables S7). The CID-4 model was the best fit for flower symmetry, indicating no relation to increased lineage diversification with the transition to vascular variants and zygomorphic flowers (*SI Appendix,* Tables S8-S9). Nevertheless, when averaging across models with AICw > 0, we found that diversification rates in lineages with zygomorphic flowers are significantly higher (p < 0.001) than those with actinomorphic flowers (Fig. 4D), suggesting some uncertainty in model selection and a potential link of this trait with the diversification process. Taken together, our results suggest a synnovation––the combined effect of the climbing growth form, tendrils, and zygomorphic flowers––not vascular variants––may be responsible for the evolutionary success of Paullinieae.

Gianoli (2) hypothesis that the climbing habit promotes species diversification in flowering plants has also been tested in the pantropical Annonaceae (magnoliids) (30) and the neotropical Nyctaginaceae (eudicots, Caryophyllales) (36), where the HiSSE model was also favored, thus corroborating our result that the acquisition of climbing is not sufficient to explain species diversification in different groups. Furthermore, Sperotto et al. (39) found diversification rates do not differ among lineages with different climbing mechanisms (e.g., tendrils vs. other mechanisms), suggesting that climbing mechanisms alone do not explain disparities in species diversification either. The last pillar of the synnovation––flower zygomorphism––is of particular note in Paullinieae. Compared to closely related genera, their zygomorphic flowers feature elaborate petal appendages along with a fleshy, bright yellow crest (16, 40), which may be an adaptive advantage by increasing flower attractiveness to pollinators or even serving as a nectar guide (40–42). Over two years of field observation of five specimens of *S. fuscifolia* revealed that the flowers were visited by 50 different insect species (41). Therefore, the link between flower types and speciation in Paullinieae may be more directly connected to specific innovations in flower structure related to pollination systems, rather than just their symmetry.

## Conclusion

In this study, we develop a robust molecular phylogeny of the Paullinieae as a framework to address fundamental questions about the evolutionary lability and developmental flexibility of their vascular system. Our findings demonstrate rampant convergent evolution of variant vascular oddities, with the rate of evolutionary transitions to variant patterns mirroring their relative developmental complexity. Given the unique suite of potential innovative traits exhibited in Paullinieae, we leveraged our datasets to discover that a potential synnovation, comprised of the acquisition of the climbing habit, tendrils, and zygomorphic flowers, collectively worked together in the diversification of this important neotropical lineage.

## Methods

### Sampling

We sampled 241 species across 333 samples, 31.83% from silica–dried specimens and 64.56% from herbarium vouchers from the United States National Herbarium (US) at the Smithsonian National Museum of Natural History (see *SI Appendix,* Table S3, Methods–– Sampling). Species names were curated by the author and Paullinieae taxonomic specialist, Dr. Pedro Acevedo–Rodríguez.

### DNA Extraction and Quality Control

DNA extractions were performed using a CTAB procedure (43) by an Autogen 965 at the Smithsonian Institution Laboratory of Analytical Biology; silica and herbarium specimens were processed separately. To maximize DNA yield, samples were divided after incubation with a lysis buffer, and two independent extractions were performed for each sample, with the extracts combined before quantification. After quantification, samples were visualized using the TapeStation 4200 (Agilent) platform to determine the molecular weight of each extraction. For each sample, 400ng of DNA was sent to Daicel Arbor Biosciences (Ann Arbor, Michigan, USA) for further quality control assessment, library preparation, target capture, and sequencing (see *SI Appendix,* Methods––DNA analysis).

### Library Preparation, Sequencing, and Data Cleaning

Samples with high, low, and mixed molecular weights were handled following different protocols that produce Illumina–compatible libraries (see *SI Appendix,* Methods–DNA analysis). The indexed libraries were quantified, and bait capture was performed following the myBaits v5.02 protocol using myBaits Expert Angiosperms– 353 baits. Samples were sequenced on the Illumina NovaSeq 6000 platform using a partial S4 PE150 lane, which yielded approximately 1 Gbps per library. The raw sequencing data underwent quality control using FastQC v0.11.9 (44). Trimmomatic v0.39 (45) was employed to remove Illumina adapters and filter out low–quality reads.

### Sequence Assembly

Consensus low–copy nuclear loci were assembled using iterative–based reference assembly with HybPiper v2.1.6 (46), with additional modification to enhance gene recovery (*SI Appendix,* Methods––DNA analysis). We filtered low–performing samples and loci to retain only those with at least 400MB of initial raw sequence data. Paralogous and putatively chimeric sequences were identified and removed (*SI Appendix,* Methods––DNA analysis). The resulting gene assemblies were manually checked in Geneious Prime v.10.11 (https://www.geneious.com). Filtered data matrices containing exons only were aligned using MAFFT v7.475 (47) and trimmed with TrimAl v1.4.1 (48) to remove sites exceeding a gap threshold of 85%.

### Phylogenetic Inference

Using the final dataset, which comprised 333 samples and 351 genes, we conducted a phylogenetic inference with the concatenated matrix and a partitioned analysis (with each gene treated as a separate partition with GTR substitution model). Maximum–likelihood tree inference was performed with IQ–TREE v1.6.12 (18) and 1000 bootstrap replicates. The input trees for ASTRAL analysis were 351 individual maximum likelihood gene trees made with IQ–TREE and 1000 bootstrap replicates under the GTR substitution model. The IQ-TREE and ASTRAL trees were compared using the cophyloplot function in the *ape* package (49) in R.

### Time–Calibrated Phylogeny

To generate the chronogram, we used the IQ–TREE consensus tree file as a topological constraint to generate 1000 bootstrap trees with varying branch lengths. We randomly sampled 100 of those bootstrap trees, containing all tips in the IQ-TREE, except three outgroups (*SI Appendix,* Methods––DNA analysis). Each of the 100 trees was pruned down to a single sample per species (including outgroups). All trees were time–calibrated using treePL (50) (see parameters in *SI Appendix,* Methods––DNA analysis). The summary result of the 100 chronograms was visualized as a maximum clade credibility tree using TreeAnnotator (51). Minimum and maximum dates were assigned to three nodes (Fig. 2), including one primary and two secondary calibrations (*SI Appendix,* Methods––DNA analysis).

### Plant Micro Technique

Specimens for developmental anatomy were obtained from three primary sources: liquid–preserved specimens collected in the field, wood collections, and herbaria(see *SI Appendix,* Tables S2, S5, and S9). Specimens were sectioned following different histological procedures to generate microscope slides for light microscopy (see *SI Appendix,* Methods––Plant Micro technique).

### Character Coding of Vascular Variants and Related Traits

To investigate the evolution of vascular variants, we characterized stem development and morpho–anatomical variations related to stem shape and eustele types (in primary growth). Information on stem development was obtained from macro– and microscopic analyses (see *SI Appendix,* Methods––Plant Micro technique) and complemented with data from the literature. We scored the stem development and the presence or absence of vascular variants for all species of Paullinieae, except *Serjania hispida*, as well as outgroups (*SI Appendix,* Table S4). For each species with vascular variants, we also specified the patterns and categories to which they belong, considering current and new anatomical information (*SI Appendix,* Methods––Character coding). Both categories (three) and patterns (six) were treated as discrete characters, as in previous studies using this system (13–15), and were utilized in phylogenetic comparative methods (e.g., stochastic character mapping).

To investigate whether the shape of young stems and the arrangement of vascular bundles are correlated, we measured the circularity of these two parameters for a dataset of 71 species included in our phylogeny. Circularity measurements, represented by fitting a circle within the outline of a specific vector object, were obtained from images of stem cross–sections at early developmental stages (e.g., primary growth or the transition from primary to secondary growth) (*SI Appendix,* Methods––Circularity). The circularity values were used to create a threshold to categorize the range of stem and eustele circularity into two groups. Stem circularity was divided into circular vs. lobed, meaning that the stem shape is cylindrical or formed by lobes and furrows, respectively. In contrast, eustele circularity was divided into typical vs. atypical, meaning that the vascular system is formed by a single ring of bundles or bundles irregularly distributed along the stem circumference, respectively. The circularity dataset was used for correlation analyses (Pearson’s correlation coefficient) among the two parameters, stochastic character mapping and test of correlated evolution between each parameter and the presence of vascular variants (see Phylogenetic Comparative Methods below).

To investigate hypotheses on the evolution of the number of peripheral vascular cylinders in species with compound stems, we focused on the genus *Serjania,* specifically the Platycoccus section, for which anatomical data were available for all nine species forming a monophyletic clade in our phylogeny. We removed from this analysis the species *S. laurotteana,* which was traditionally included in this section (52), as it is distantly related to the remaining species of the group. To understand the diversity in peripheral cylinder numbers, we counted vascular bundles (mostly by identifying protoxylem poles) and the number of peripheral vascular cylinders in adult stems. This data was scored for all species of the Platycoccus section and used to inform the hypothesis of the ancestral state of the section.

### Phylogenetic Comparative Methods

To infer the evolutionary history of vascular variants and other morphological traits in Paullinieae, we leveraged our anatomical database (*SI Appendix,* Table S4) to match information on vascular variants for 218 of the 224 species in our chronogram (excluding six names, three identified as “sp.” and three outgroups not belonging to the supertribe Paulliniodae). The same dataset was used to illustrate the section classification across Paullinieae and trait–dependent diversification analyses (see below). The evolution of stem and eustele circularity (*SI Appendix,* Table S5) was examined in a reduced dataset (including 71 species), trimmed from the 218 species–vascular variants dataset. Ancestral trait estimates for vascular variants, stem/eustele circularity, and bundle count were inferred using stochastic mapping as implemented in the R package “phytools” (53). The best model was selected based on the AIC scores of different evolution models, including equal rates, symmetric rates, and all rates different. One thousand stochastic character maps were simulated to estimate the posterior probabilities of the ancestral states for each character, and the results were summarized using the “make.simmap” function in *phytools* (53). Before ancestral state estimates, the phylogenetic signal of vascular variants was estimated using Pagel’s λ (lambda) with the “fit.discrete” function in *phytools* (53). For vascular variants, we inferred the ancestral states of the three discrete *categories* and six *patterns* of vascular variants in two independent analyses (*SI Appendix,* Methods—Character coding). In addition to stochastic mapping, we also used “PARAMO” (Phylogenetic Ancestral Reconstruction of Anatomy by Mapping Ontologies) (54) to infer the evolution of vascular variants. PARAMO reconstructed the evolution of vascular variants using hidden, structured Markov models and stochastic mapping, enabling the modeling of interdependencies between different developmental events that lead to the formation of similar complex traits (*SI Appendix*, Methods––Character coding). For the test of correlated evolution between stem circularity and eustele circularity with the emergence of vascular variants, we used Pagel’s 1994 phylogenetic test, as implemented in the “fit.pagel” function in *phytools*. To test whether vascular variants (discrete trait) are influenced by stem and vascular circularity (continuous trait), we fitted the threshold model for the same circularity dataset using the “threshBayes” function in *phytools.* All analyses were performed using the R software (55).

### Diversification Rates

To infer shifts in diversification rates, we applied BAMM (56) and MEDUSA(57) to a chronogram pruned to include only one individual per species. We provided a complete list of species and calculated the proportion of species sampled per genus, which was used to generate the richness file for MEDUSA and define the sampling fractions for BAMM. For BAMM, priors were generated using the R package BAMMtools (58) for four independent MCMC chains run for 10M generations, sampling every 1000 generations. Convergence was assessed using CODA (59). Rates-through-time plots were also generated for speciation (λ), extinction (μ), and net diversification (r) using “PlotRateThroughTime” for Paulliniodeae.” For MEDUSA, we also accounted for incomplete taxon sampling and tested for mixed models (default), in which parameter estimates are weighted means across the other two models: one that considers speciation only and the other that considers both speciation and extinction. All analyses were performed using R software (55).

### Trait–Dependent Diversification

To test whether vascular variants, liana habit, tendrils, and zygomorphic flowers are associated with speciation and diversification rate shifts, we applied diverse models with Hidden State Speciation and Extinction (60). HiSSE uses a hidden Markov model to account for an unobserved trait that may interact with the observed one. The hidden states of the unobserved character can influence diversification rates on their own or together with the observed trait (*SI Appendix*, Table S7). Here, for each trait, we compared character–dependent (e.g., BiSSE-like, HiSSE) and character–independent (e.g., CID-2 and CID-4) models of diversification, contrasting with a null model containing equal turnover and extinction fractions and no hidden character (*SI Appendix*, Table S8). We accounted for incomplete taxon sampling by specifying sampling fractions for each character state (SI Appendix, Table S8). Model selection was based on Akaike Information Criterion (AIC) and Akaike weights (AICw) to identify the best-fitting model. All analyses were performed using the R software (55).

## Data, Materials, and Software Availability

Raw reads have been deposited in the NCBI Sequence Read Archive under SUB15448216, trees and alignments are deposited at https://zenodo.org/records/15867052, and scripts and associated datasets are deposited at https://github.com/joycechery/paullinieae_phylogenomics. All other study data are included in the article and/or SI Appendix.

## Acknowledgments

We are grateful to those who contributed leaf and stem samples over the years dedicated to studying Paullinieae lianas, including the NY and US herbaria, for allowing the destructive sampling of multiple specimens. We would also like to thank Gabriel Johnson, who carried out DNA extractions and cleaning at the Smithsonian Lab of Analytical Biology, and Dr. Kristin Winchell for providing invaluable feedback on the draft manuscript. This work was supported in part through the NYU IT High Performance Computing resources, services, and staff expertise. This work was funded by startup grants to Cornell University and New York University to J.G.O. and an NSF #2401675 to J.G.O.

## References

1. S. Isnard, W. K. Silk, Moving with climbing plants from Charles Darwin’s time into the 21st century. American J of Botany 96, 1205–1221 (2009).

2. E. Gianoli, Evolution of a climbing habit promotes diversification in flowering plants. Proceedings of the Royal Society of London. Series B: Biological Sciences 271, 2011–2015 (2004).

3. E. Gianoli, The behavioural ecology of climbing plants. AoB PLANTS 7 (2015).

4. G. M. F. van der Heijden, J. S. Powers, S. A. Schnitzer, Lianas reduce carbon accumulation and storage in tropical forests. Proceedings of the National Academy of Sciences 112, 13267–13271 (2015).

5. S. Estrada-Villegas, S. S. Pedraza Narvaez, A. Sanchez, S. A. Schnitzer, Lianas Significantly Reduce Tree Performance and Biomass Accumulation Across Tropical Forests: A Global Meta-Analysis. Front. For. Glob. Change 4, 812066 (2022).

6. V. Angyalossy, G. Angeles, M. R. Pace, A. C. Lima, “Liana anatomy: a broad perspective on structural evolution of the vascular system.” in Ecology of Lianas., (John Wiley & Sons, Ltd, 2015), pp. 253–287.

7. J. G. Onyenedum, M. R. Pace, The role of ontogeny in wood diversity and evolution. American Journal of Botany 108, 2331–2355 (2021).

8. I. L. Cunha Neto, Vascular variants in seed plants—a developmental perspective. AoB PLANTS 15, 1–15 (2023).

9. N. P. Rowe, T. Speck, “The evolution of angiosperm lianescence: a perspective from xylem structure-function.” in Ecology of Lianas., (JohnWiley & Sons, 2015), pp. 221–250.

10. H. Schenck, “Beiträge zur Biologie und Anatomie der Lianen im Besonderen der in Brasilien einheimischen Arten.” in Beiträge Zur Anatomie Der Lianen: Botanische Mittheilungen Aus Den Tropen., (Verlag von Gustav Fisher., 1893).

11. L. Radlkofer, “Sapindaceae” in Das Planzenreich IV, 165 (Heft 98a-h), (W. Engelmann, 1931-1933), pp. 1–1539.

12. M. R. Pace, et al., The wood anatomy of Sapindales: diversity and evolution of wood characters. Braz. J. Bot 45, 283–340 (2022).

13. J. G. Chery, M. R. Pace, P. Acevedo-Rodríguez, C. D. Specht, C. J. Rothfels, Modifications during Early Plant Development Promote the Evolution of Nature’s Most Complex Woods. Current Biology 30, 237-244.e2 (2020).

14. I. L. Cunha Neto, et al., Molecular phylogeny of Urvillea (Paullinieae, Sapindaceae) and its implications in stem vascular diversity. Annals of Botany mcad093 (2023). 10.1093/aob/mcad093.

15. N. F. Marques, I. L. Cunha Neto, L. A. Brito, G. V. Somner, Serjania piscatoria (Paullinieae, Sapindaceae) as a symbol of vascular variants polymorphism. Botanical Journal of the Linnean Society 208, 58–74 (2025).

16. P. Acevedo-Rodríguez, et al., Generic Relationships and Classification of Tribe Paullinieae (Sapindaceae) with a New Concept of Supertribe Paulliniodae. sbot 42, 96–114 (2017).

17. M. G. Johnson, et al., A Universal Probe Set for Targeted Sequencing of 353 Nuclear Genes from Any Flowering Plant Designed Using k-Medoids Clustering. Syst Biol 68, 594– 606 (2019).

18. L.-T. Nguyen, H. A. Schmidt, A. von Haeseler, B. Q. Minh, IQ-TREE: A Fast and Effective Stochastic Algorithm for Estimating Maximum-Likelihood Phylogenies. Molecular Biology and Evolution 32, 268–274 (2015).

19. S. Mirarab, et al., ASTRAL: genome-scale coalescent-based species tree estimation. Bioinformatics 30, i541–548 (2014).

20. J. D. Urdampilleta, E. R. Forni-Martins, M. S. Ferrucci, Phylogenetics of the supertribe Paulliniodae (Sapindaceae) with emphasis on chromosome evolution: Taxonomic implications including the new genus Acevedoa. Annals of Botany mcaf049 (2025). 10.1093/aob/mcaf049.

21. J. G. Chery, P. Acevedo-Rodríguez, C. J. Rothfels, C. D. Specht, Phylogeny of Paullinia L. (Paullinieae: Sapindaceae), a diverse genus of lianas with dynamic fruit evolution. Molecular Phylogenetics and Evolution 140, 106577 (2019).

22. V. W. Steinmann, M. S. Ferrucci, C. A. Maya-Lastra, Phylogenetics of Serjania (Sapindaceae-Paullinieae), with emphasis on fruit evolution and the description of a new species from Michoacán, Mexico. Systematics and Biodiversity 20, 1–21 (2022).

23. H. Medeiros, P. Acevedo-Rodríguez, J. de Carvalho Lopes, R. C. Forzza, Taxonomic revision and phylogenetic relationships of Thinouia (Sapindaceae), a neotropical genus. PhytoKeys 252, 207–273 (2025).

24. Flora e Funga do Brasil, Jardim Botânico do Rio de Janeiro. Available at: http://floradobrasil.jbrj.gov.br/ [Accessed 14 November 2024].

25. S. Buerki, et al., An updated infra‐familial classification of Sapindaceae based on targeted enrichment data. Am J Bot 108, 1234–1251 (2021).

26. E. M. Joyce, et al., Phylogenomic analyses of Sapindales support new family relationships, rapid Mid-Cretaceous Hothouse diversification, and heterogeneous histories of gene duplication. Front. Plant Sci. 14 (2023).

27. S. Buerki, F. Forest, T. Stadler, N. Alvarez, The abrupt climate change at the Eocene– Oligocene boundary and the emergence of South-East Asia triggered the spread of sapindaceous lineages. Ann Bot 112, 151–160 (2013).

28. M. Lu, M. Fradera-Soler, F. Forest, T. G. Barraclough, O. M. Grace, Evidence linking life-form to a major shift in diversification rate in Crassula. American Journal of Botany 109, 272–290 (2022).

29. J. Chave, et al., Evidence for a Miocene pulse of diversification of the tropical American clade of the Brazil nut family (Lecythidaceae). Botany Letters 171, 537–551 (2024).

30. B. Xue, et al., Accelerated diversification correlated with functional traits shapes extant diversity of the early divergent angiosperm family Annonaceae. Molecular Phylogenetics and Evolution 142, 106659 (2020).

31. C. Hoorn, L. G. Lohmann, L. M. Boschman, F. L. Condamine, Neogene History of the Amazonian Flora: A Perspective Based on Geological, Palynological, and Molecular Phylogenetic Data. Annual Review of Earth and Planetary Sciences 51, 419–446 (2023).

32. M. R. Pace, L. G. Lohmann, V. Angyalossy, The rise and evolution of the cambial variant in Bignonieae (Bignoniaceae). Evolution and Development 11, 465–479 (2009).

33. L. Luna-Márquez, W. V. Sharber, B. A. Whitlock, M. R. Pace, Ontogeny, anatomical structure and function of lobed stems in the evolution of the climbing growth form in Malvaceae (Byttneria Loefl.). Annals of Botany 128, 859–874 (2021).

34. I. L. Cunha Neto, J. G. Onyenedum, Ectopic cambia: Connections between natural and experimental vascular mutants. American J of Botany 110, e16246 (2023).

35. I. L. Cunha Neto, A. A. Snead, J. B. Landis, C. D. Specht, J. G. Onyenedum, Ectopic cambia in Japanese wisteria (Wisteria floribunda) vines are associated with the expression of conserved KNOX genes. [Preprint] (2024). Available at: http://biorxiv.org/lookup/doi/10.1101/2024.08.07.606835 [Accessed 13 August 2024].

36. I. L. Cunha Neto, M. R. Pace, R. Hernández-Gutiérrez, V. Angyalossy, Linking the evolution of development of stem vascular system in Nyctaginaceae and its correlation to habit and species diversification. EvoDevo 13, 4 (2022).

37. I. L. Cunha Neto, et al., Medullary bundles in Caryophyllales: form, function, and evolution. New Phytologist nph.19342 (2023). 10.1111/nph.19342.

38. A. H. Gentry, “The distribution and evolution of climbing plants” in The Biology of Vines, (Cambridge University Press, 1991), pp. 3–49.

39. P. Sperotto, N. Roque, P. Acevedo-Rodríguez, T. Vasconcelos, Climbing mechanisms and the diversification of neotropical climbing plants across time and space. New Phytologist 240, 1561–1573 (2023).

40. H. A. de Lima, G. V. Somner, A. M. Giulietti, Duodichogamy and sex lability in Sapindaceae: the case of Paullinia weinmanniifolia. Plant Syst Evol 302, 109–120 (2016).

41. M. Souza, M. Milaneze-Gutierre, L. Souza, Flower structure of Serjania fuscifolia (Sapindaceae) and its relationship with visiting insect. International Journal of Life Science Research Archive 03, 62–074 (2022).

42. C. dos Santos Almeida, G. V. Somner, B. de Sá-Haiad, Floral anatomy in Serjania clematidifolia (Paullinieae, Sapindaceae): Insights into a monoecious sexual system with multicyclic dichogamy. Flora 320, 152614 (2024).

43. J. Doyle, “DNA Protocols for Plants” in Molecular Techniques in Taxonomy, G. M. Hewitt, A.W. B. Johnston, J. P. W. Young, Eds. (Springer, 1991), pp. 283–293.

44. S. Andrews, FastQC: A Quality Control Tool for High Throughput Sequence Data [Online]. (2010). Available at: http://www.bioinformatics.babraham.ac.uk/projects/fastqc/.

45. A. M. Bolger, M. Lohse, B. Usadel, Trimmomatic: a flexible trimmer for Illumina sequence data. Bioinformatics 30, 2114–2120 (2014).

46. M. G. Johnson, et al., HybPiper: Extracting coding sequence and introns for phylogenetics from high-throughput sequencing reads using target enrichment. Applications in Plant Sciences 4, apps.1600016 (2016).

47. K. Katoh, D. M. Standley, MAFFT Multiple Sequence Alignment Software Version 7: Improvements in Performance and Usability. Molecular Biology and Evolution 30, 772–780 (2013).

48. S. Capella-Gutiérrez, J. M. Silla-Martínez, T. Gabaldón, trimAl: a tool for automated alignment trimming in large-scale phylogenetic analyses. Bioinformatics 25, 1972–1973 (2009).

49. E. Paradis, K. Schliep, ape 5.0: an environment for modern phylogenetics and evolutionary analyses in R. Bioinformatics 35, 526–528 (2019).

50. S. A. Smith, B. C. O’Meara, treePL: divergence time estimation using penalized likelihood for large phylogenies. Bioinformatics 28, 2689–2690 (2012).

51. A. J. Drummond, A. Rambaut, BEAST: Bayesian evolutionary analysis by sampling trees. BMC Evolutionary Biology 7, 214 (2007).

52. P. Acevedo-Rodríguez, “Systematics of Serjania (Sapindaceae). Part I: A revision of Serjania Sect. Platycoccus. Doctoral dissertation - New York Botanical Garden,” New York Botanical Garden. (1993).

53. L. J. Revell, phytools: an R package for phylogenetic comparative biology (and other things). Methods Ecol Evol 3, 217–223 (2012).

54. S. Tarasov, I. Mikó, M. J. Yoder, J. C. Uyeda, PARAMO: A Pipeline for Reconstructing Ancestral Anatomies Using Ontologies and Stochastic Mapping. Insect Systematics and Diversity 3, 1 (2019).

55. R Core Team, R: A Language and Environment for Statistical Computing. (2017). Deposited 2017.

56. D. L. Rabosky, Automatic Detection of Key Innovations, Rate Shifts, and Diversity-Dependence on Phylogenetic Trees. PLOS ONE 9, e89543 (2014).

57. M. E. Alfaro, et al., Nine exceptional radiations plus high turnover explain species diversity in jawed vertebrates. Proc Natl Acad Sci U S A 106, 13410–13414 (2009).

58. D. L. Rabosky, et al., BAMMtools: an R package for the analysis of evolutionary dynamics on phylogenetic trees. Methods in Ecology and Evolution 5, 701–707 (2014).

59. M. Plummer, N. Best, K. Cowles, CODA: Convergence Diagnosis and Output Analysis for MCMC. 6 (2006).

60. J. M. Beaulieu, B. C. O’Meara, Detecting Hidden Diversification Shifts in Models of Trait-Dependent Speciation and Extinction. Systematic Biology 65, 583–601 (2016).

61. P. Acevedo-Rodríguez, Lianas and climbing plants of the Neotropics. (2015). Available at: https://naturalhistory.si.edu/research/botany/research/lianas-and-climbing-plants-neotropics.

62. I. L. D. Cunha Neto, F. M. Martins, G. V. Somner, N. Tamaio, Successive cambia in liana stems of Paullinieae and their evolutionary significance in Sapindaceae. Botanical Journal of the Linnean Society 186, 66–88 (2018).

