## Supplemental Figs 1-12, and Tables 2,5,9 for "Rampant convergent evolution of vascular oddities and a synnovation characterize the rapid radiation of Paullinieae lianas"

PNAS

**This PDF file includes:**

Figures S1 to S12
Tables S2, S5, S6 to S9 (other tables are provided as external files)
Supplementary methods
SI References

**External files (data deposited at https://zenodo.org/records/15867052)**

**Table S1.** Accession database. Plant specimens used in the Paullinieae phylogeny, including their taxonomic classifications, collection details, and herbarium information.

**Table S3.** Sequencing and assembly statistics.

**Table S4.** Anatomical database. Detailed anatomical data for species in the Paullinieae tribe and outgroups, focusing on stem development and vascular variants. The species list included in this database also comprises the curated list of species in Paullinieae by Dr. Pedro Acevedo-Rodríguez, as well as the total number of species accepted in this study.

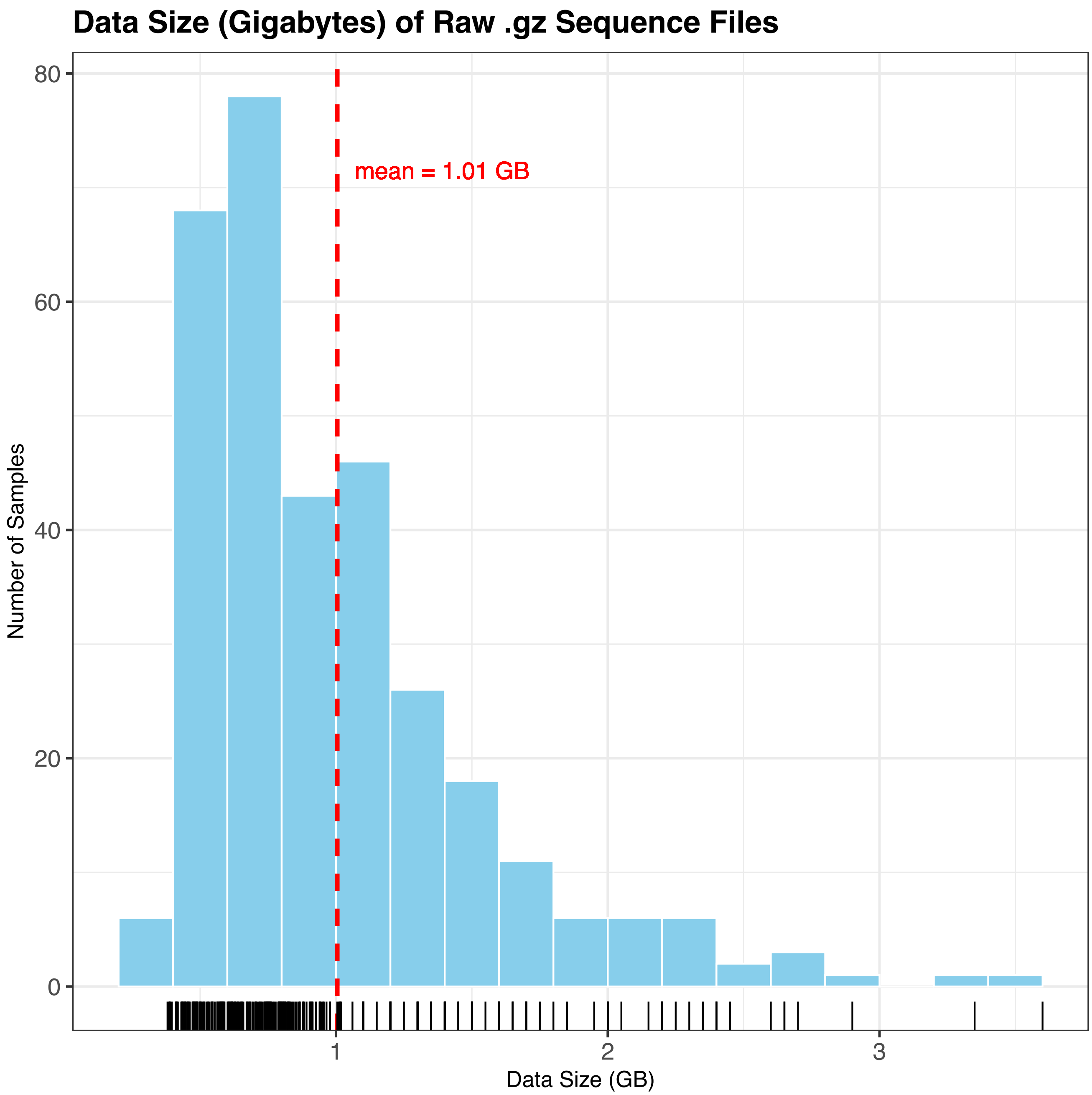


**Figure S1.** Histogram of the data size for the 321 newly sequenced samples produced in this study. This is the mean data size per sample, averaging across both R1 and R2 .gz files for a given sample. The mean data size across all samples is 1.01 GB.

PNAS

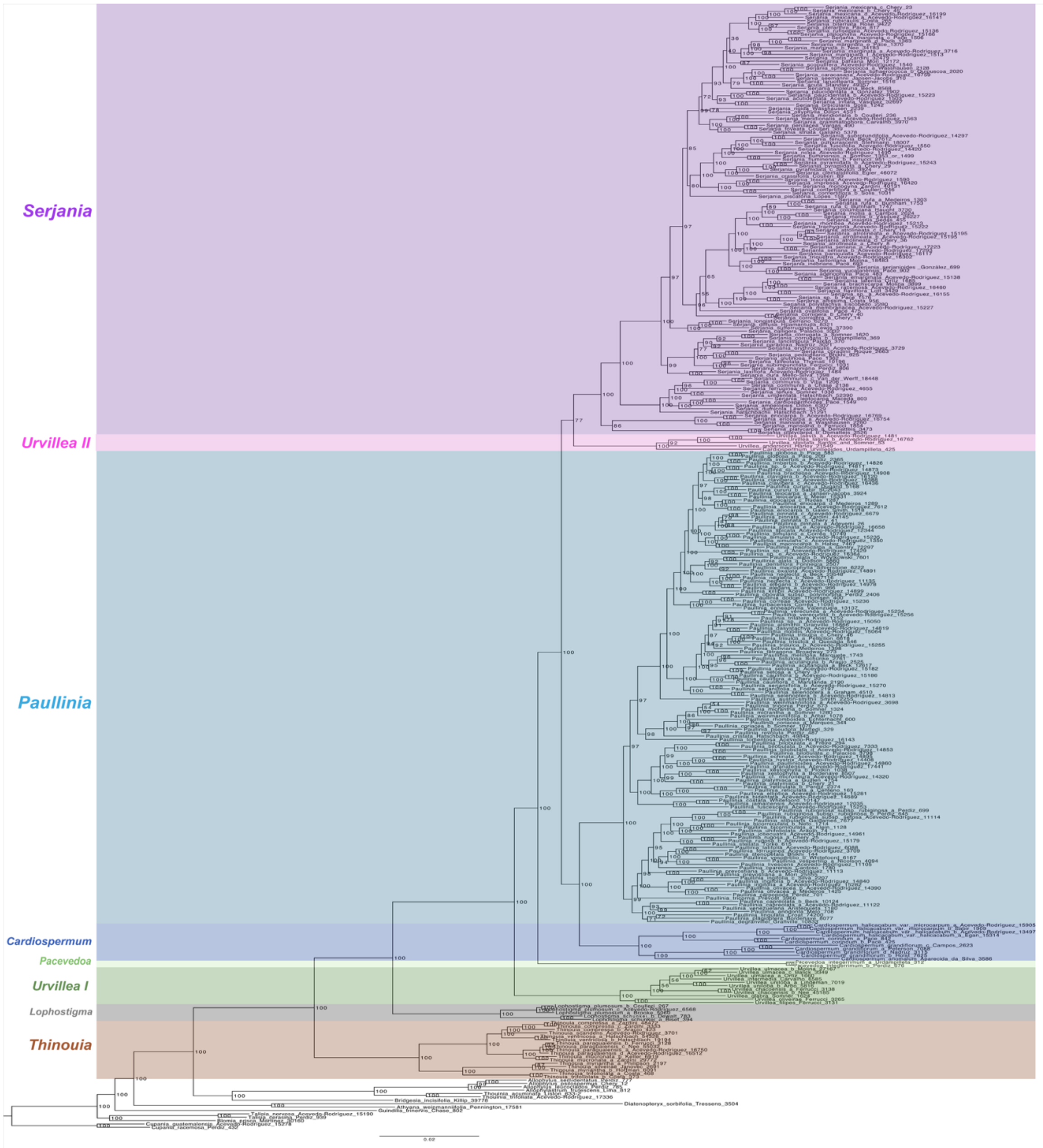


**Figure S2.** Maximum likelihood phylogeny (IQ-TREE) inferred from a partitioned concatenated analysis using 351 Angiosperm 353 nuclear exons for 212 Paullinieae species and 15 outgroups. Branches are labeled with bootstrap values. The color-coding indicates major clades of Paullinieae.

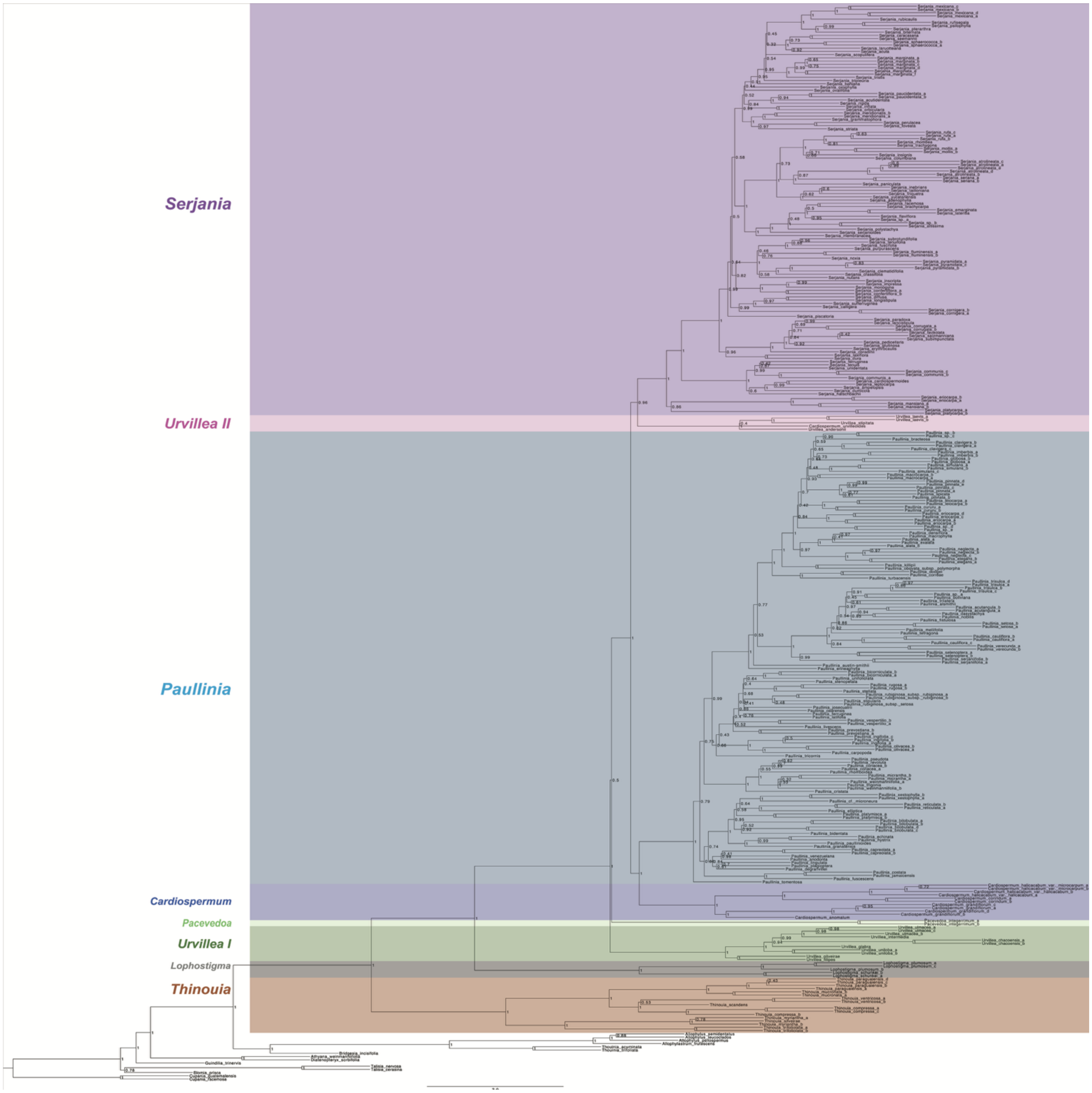


**Figure S3.** ASTRAL tree inferred from the gene tree of 351 Angiosperm 353 nuclear exons for 212 Paullinieae species and 15 outgroups. Branches are labeled with bootstrap values. The color-coding indicates major clades of Paullinieae.

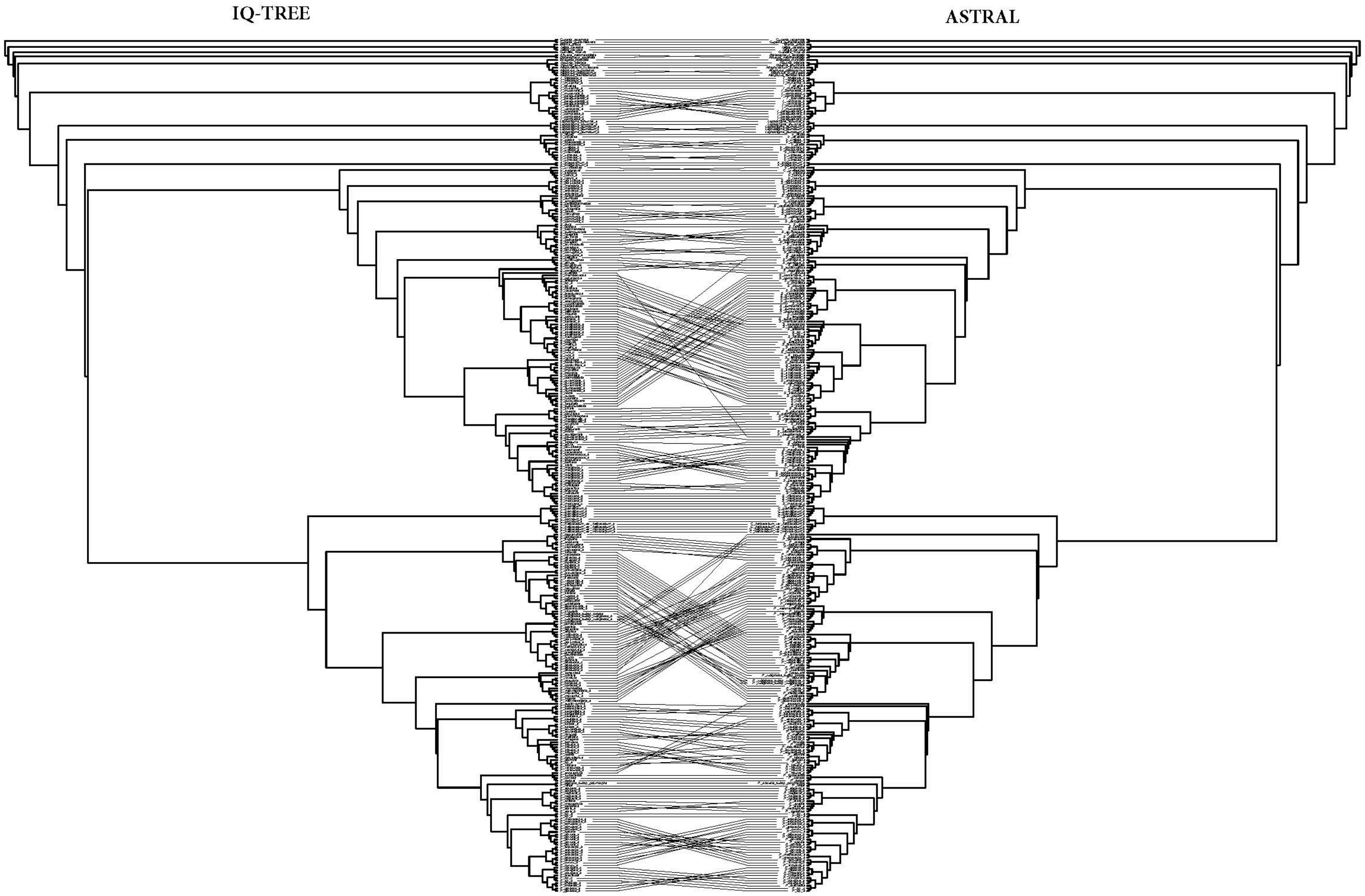


**Figure S4**. Topological comparison between IQ-TREE (left) and ASTRAL (right) tree inferences. The backbone relationships concerning the 8 major clades are consistent across trees. All conflicts are infrageneric.

PNAS

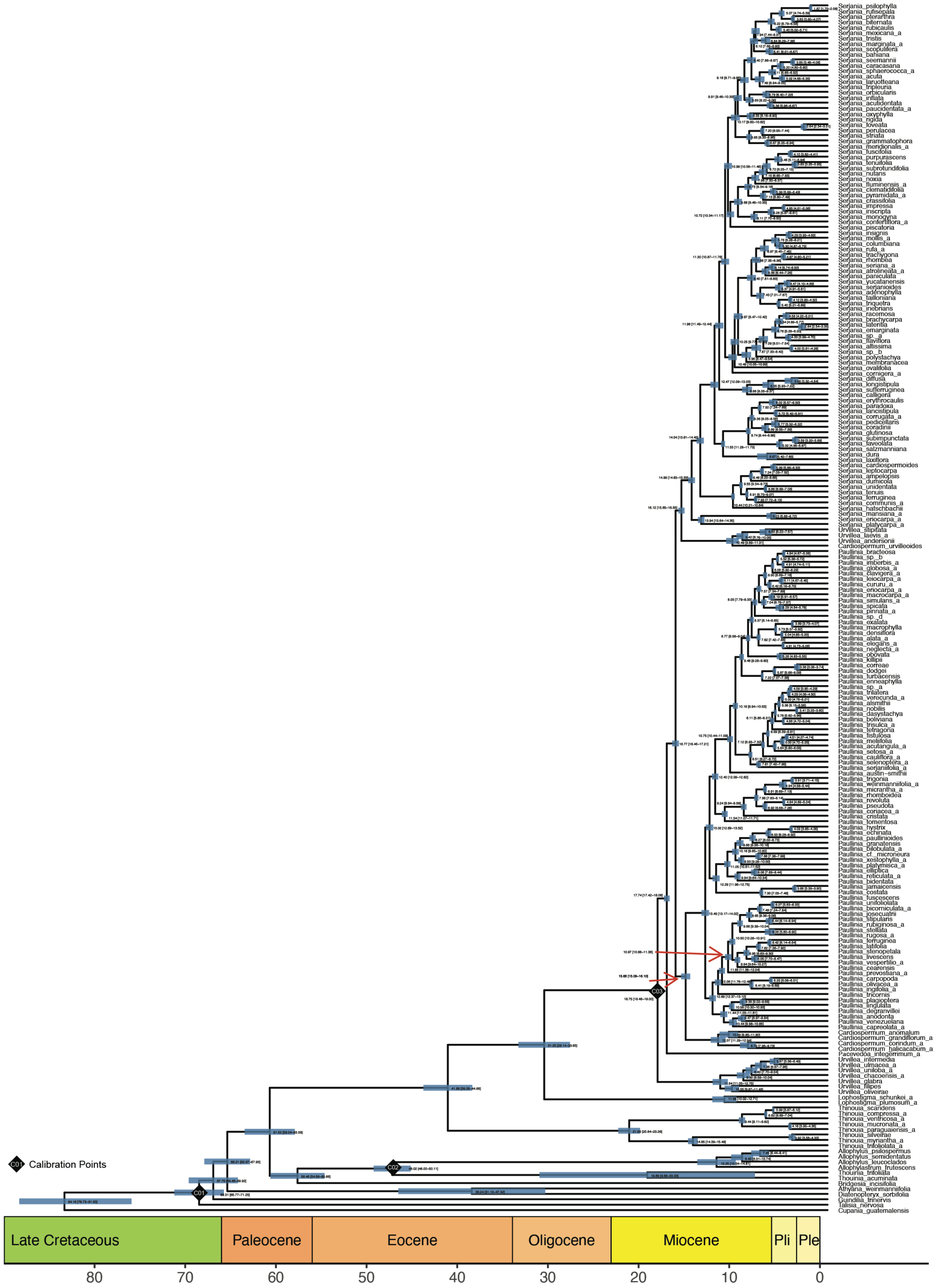


**Figure S5.** Chronogram of Paullinieae as estimated under a penalized likelihood model through TreePL based on two secondary calibrations (1) and one primary calibration based on a root fossil, *Amperilorhiza heteroxylon* (2). Blue bars represent 95% highest posterior densities (HPDs).

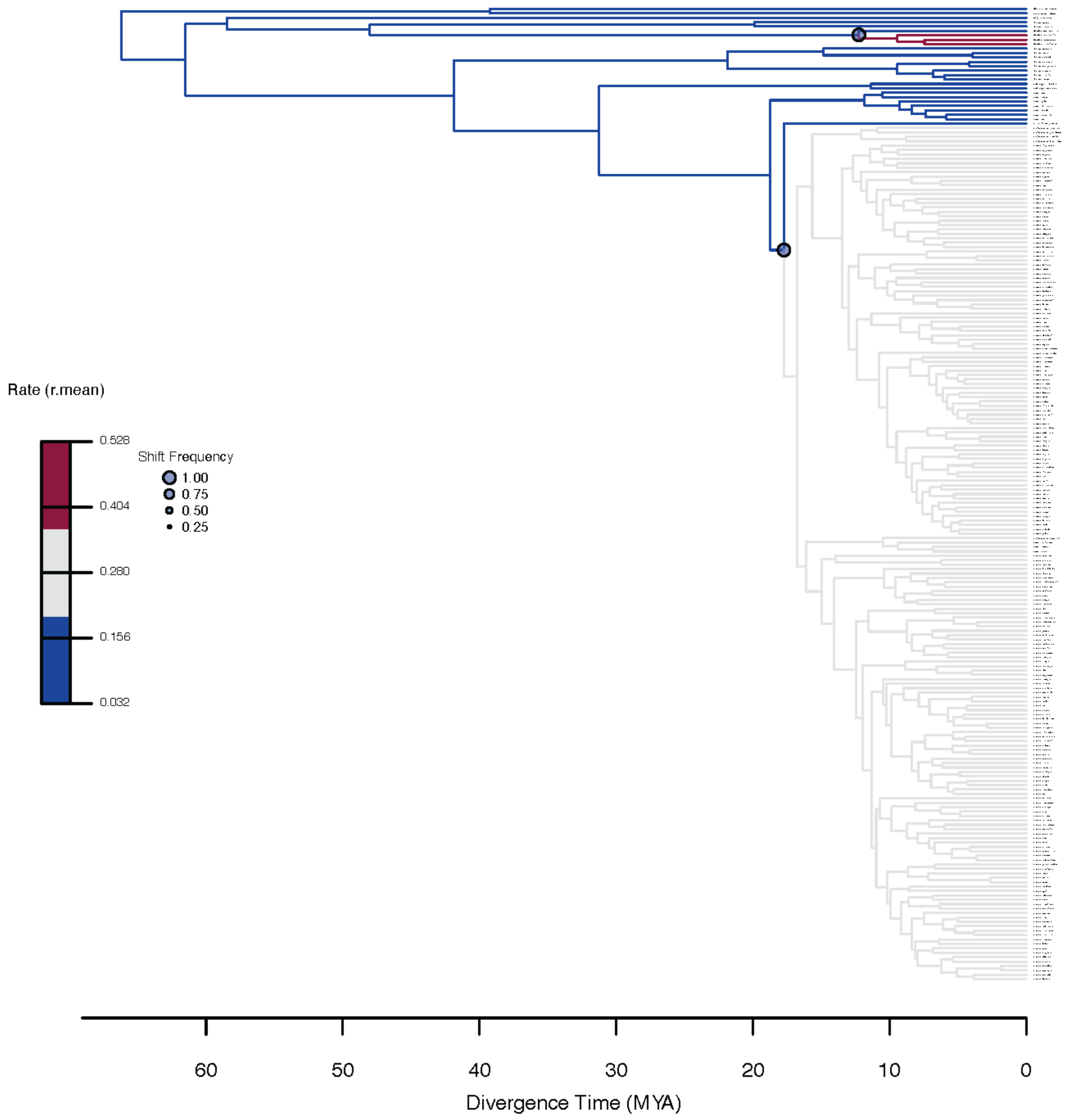


**Figure S6.** Diversification rate shifts in the Paullinieae phylogeny using MEDUSA. The net diversification rates are calculated as rates per lineage per million years. Circles indicate rate shifts, with the size of each circle representing its frequency. The unit for the geological scale is million years. Branch colors represent estimates of diversification rates.

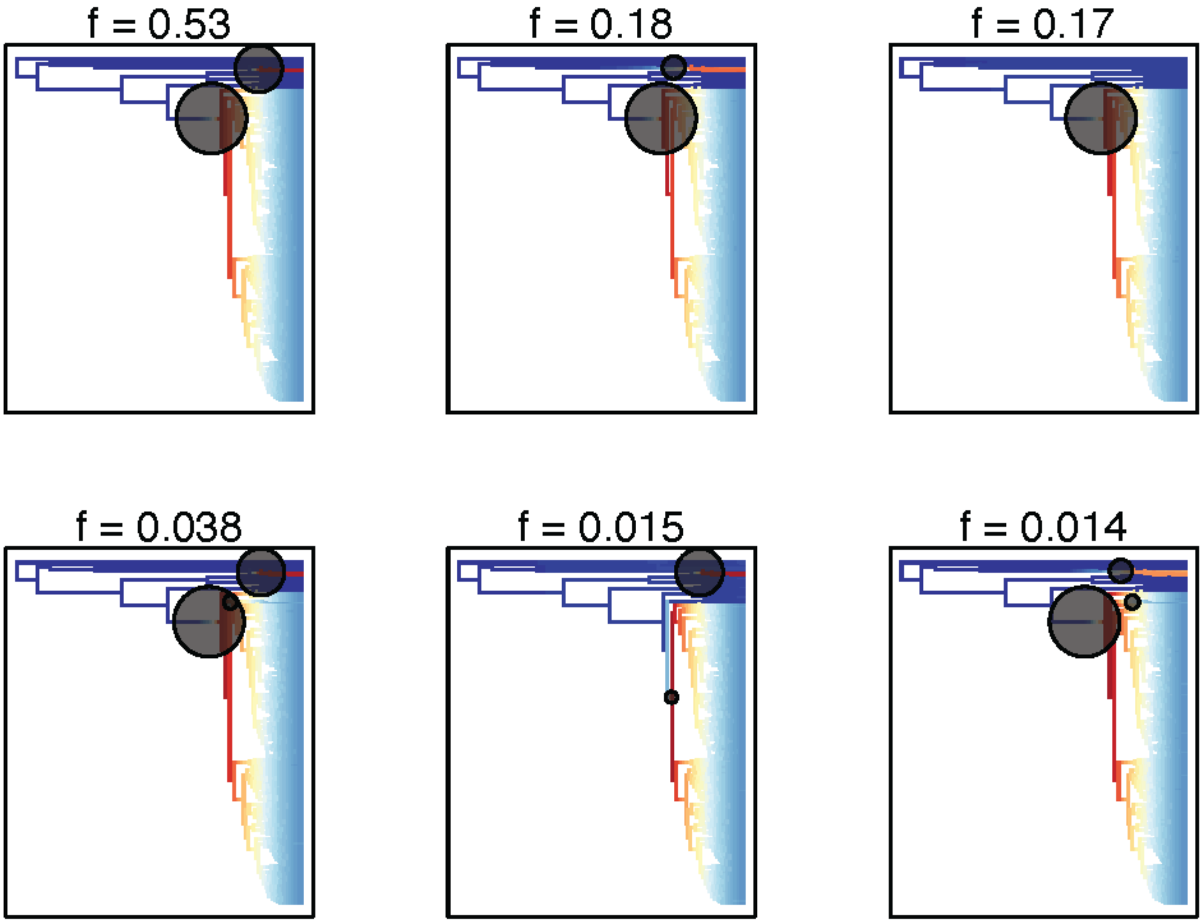


**Figure S7.** The six most credible shifts configurations detected by BAMM. In the most credible shift (f = 0.53), the shifts correspond to *Allophyllus* + *Allophylastrum* (smaller circle) and Paullinieae, excluding *Thinouia* + *Lophostigma* (bigger circle). Branch colors represent estimates of diversification rates; red indicates higher rates, while blue indicates lower rates.

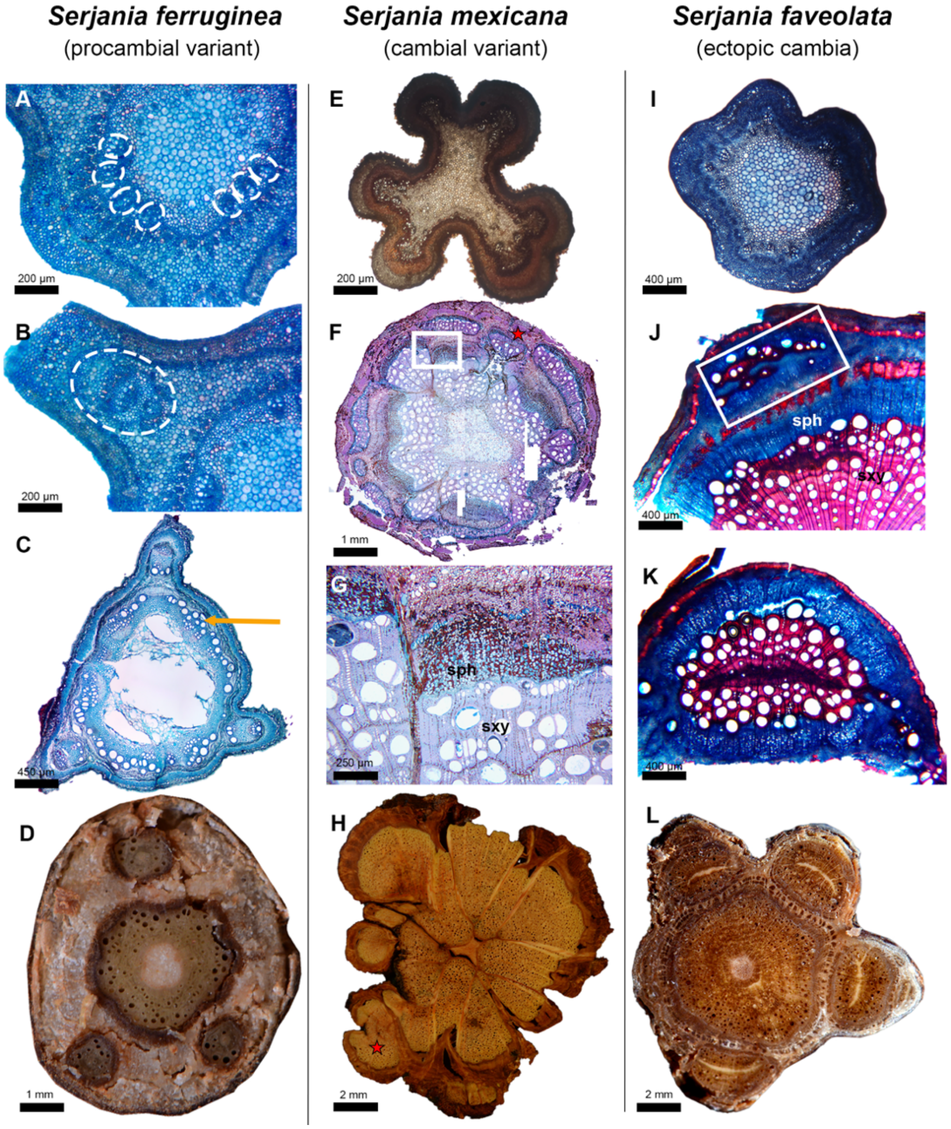


**Figure S8.** Ontogeny of vascular variants in Paullinieae lianas. All images are in cross-section. *(A–D)* Compound stem; *Serjania ferruginea. (A–B)* Primary growth characterized by a lobed shape and atypical eustele. (*A*) White dashed circles highlight the vascular bundles, which will form the central cylinder (*B*). White dashed circle highlights vascular bundles that will form the peripheral cylinder. (*C*) Early secondary growth characterized by the production of additional vascular tissues by the cambium in the central cylinder and the three peripheral cylinders. The arrow indicates early secondary xylem. (*D*) Developed stem with a central vascular cylinder and three peripheral cylinders. *(E–H)* Phloem wedges; *S. mexicana.* (*E*) Primary growth characterized by a lobed shape and atypical eustele. (*F–G*) Secondary growth showing phloem wedges (box in *F* indicates the inset in *G*), derived from an increased production of secondary phloem (sph) compared to secondary xylem (sxy) in the same cambium sector. (*F*) Note also the presence of ectopic cambia (red star), as an additional pattern. (*H*) Mature stem with phloem wedges and additional vascular units derived from ectopic cambia (red star). (*I–L*) Ectopic cambia; *S. faveolata.* (*I*) Primary growth characterized by a circular shape and a typical eustele. (*J*) Secondary growth shows the establishment of a vascular tissue derived from a new cambium (box) initiated from vascular parenchyma internally to the pericycle ring. (*K*) Developed vascular unit derived from the new (circular) cambium. (*L*) Mature stem with four well-developed peripheral vascular cylinders derived from ectopic cambia.

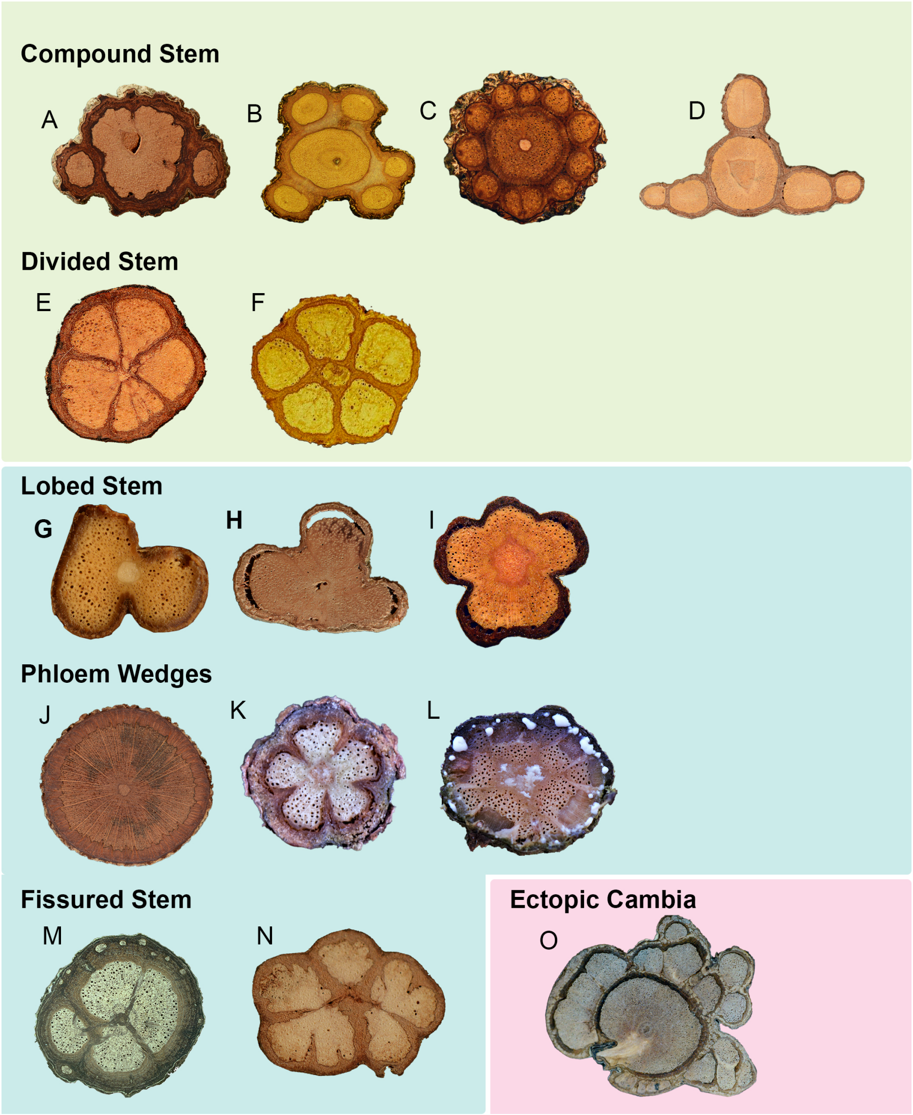


**Figure S9.** Diversity of vascular variants categories and patterns in Paullinieae lianas. (*A–E*) Procambial variants. *(A-D)* Compound stem; the number of peripheral vascular cylinders varies from two to ten. (*A*) *Serjania scopulifera.* (*B*) *Serjania glabrata.* (*C*) *Serjania clematidifolia.* (*D*) *Serjania nutans.* (*E-F*) Divided stem; usually formed by five peripheral cylinders without a central cylinder (*E*); a central cylinder may be formed late in development, which generally involves de novo cambia (*F*). (*E*) *Serjania corrugata.* (*F*) *Serjania corrugata*; reproduced from Rizzieri et al. (3), with permission from Brill). (*G-I*) Lobed stems; they can vary in the number of lobes and furrows, typically having two, three, or five. (G–O) Cambial variants. (*G*) *Urvillea chacoensis*. (*H*) *Paullinia obovata.* (*I*) *Paullinia stellata.* (*J-M*) Phloem wedges; the size and depth of the wedges are variable, ranging from many shallow wedges (*J*) to a few deep ones (*K*). Most cases contain a few relatively deep square wedges that are regularly defined by delimiting, wide rays (*L* and *M*). (*J*) *Paullinia latifolia.* (*K*) *Serjania salzmanniana.* (*L*) *Serjania rubicaulis.* (*M–N*) Fissured stems; the vascular system is dissected into a few (*M*) or multiple compartments (*N*). (*M*) *Serjania piscatoria;* note also the formation of ectopic cambia in the periphery (reproduced from Marques et al. (4), with permission from The Linnean Society of London). (*N*) *Urvillea laevis.* (*O*) Ectopic cambia; they can be formed as eccentric arcs (this figure) or concentric rings (Fig. 3), neoformations (*M*), or complete cylinders (Fig. S9). (*O*) *Paullinia pseudota.* Images are not to scale. Stem diameter ranges from 10 to 40 mm. Image credit: Pedro Acevedo-Rodríguez (5), except when stated otherwise.

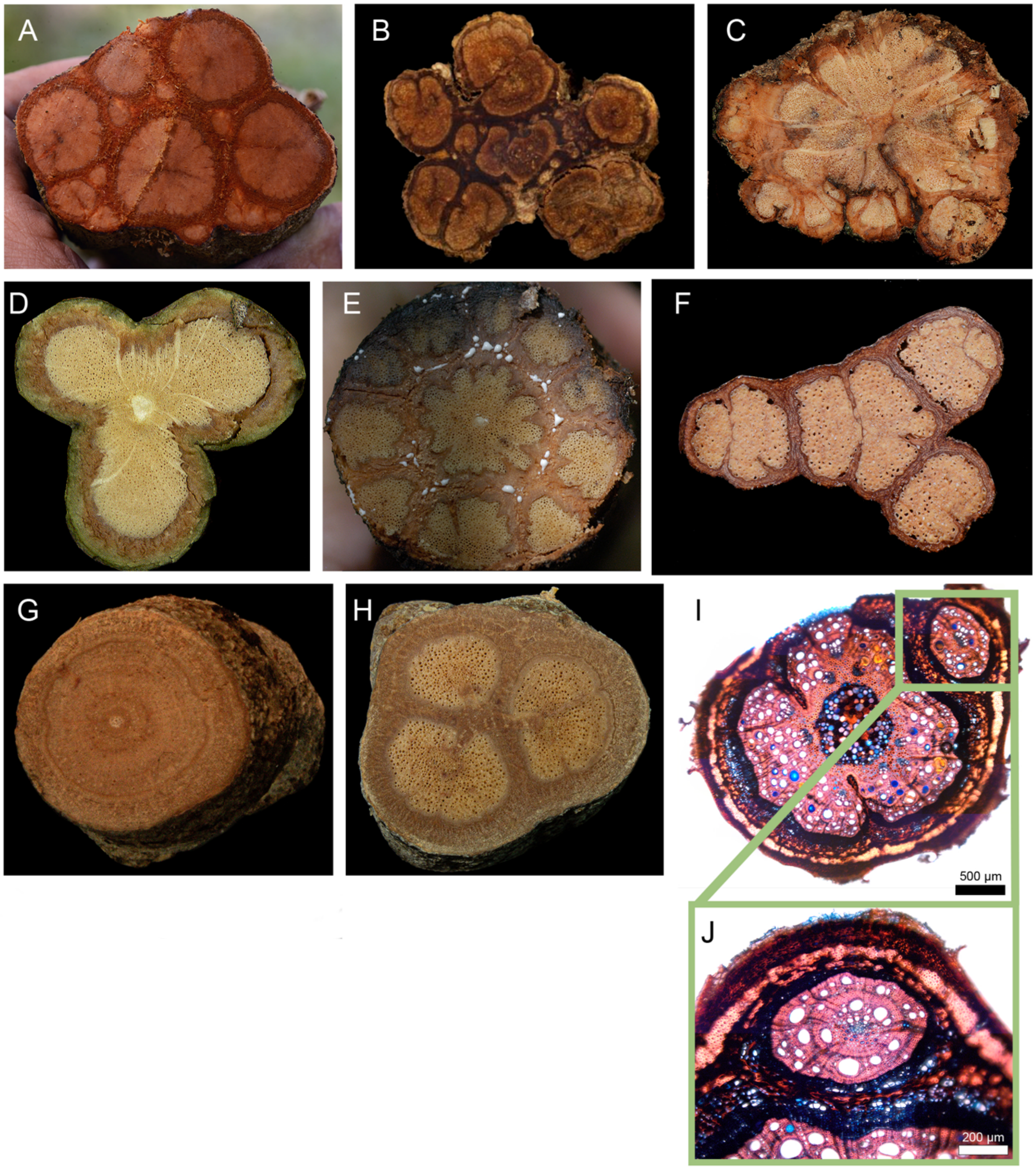


**Figure S10.** Examples of combinations of vascular variants in Paullinieae lianas. The combinations primarily include the emergence of ectopic cambia and phloem wedges/fissured stems in species with other patterns. *(A)* Compound stem + ectopic cambia; *Serjania neei.* (*B*) Divided stem + ectopic cambia; *Serjania corrugata*. *(C)* Phloem wedges + ectopic cambia; *Serjania mexicana. (D)* Lobed stem + phloem wedges; *Paullinia caloptera. (E)* Compound stem + phloem wedges; *Serjania pyramidata*. (*F*) *Serjania unguiculata*; compound stem + fissured stem. (*G-I)* Polymorphism in *S. piscatoria,* which can display typical (*G*), fissured (*H*), or compound stems (*I*). Stem diameter ranges from 10 to 50 mm. Image credit: Pedro Acevedo-Rodríguez (5), except for *(G-H)* (reproduced from Marques et al. (4), with permission from The Linnean Society of London).

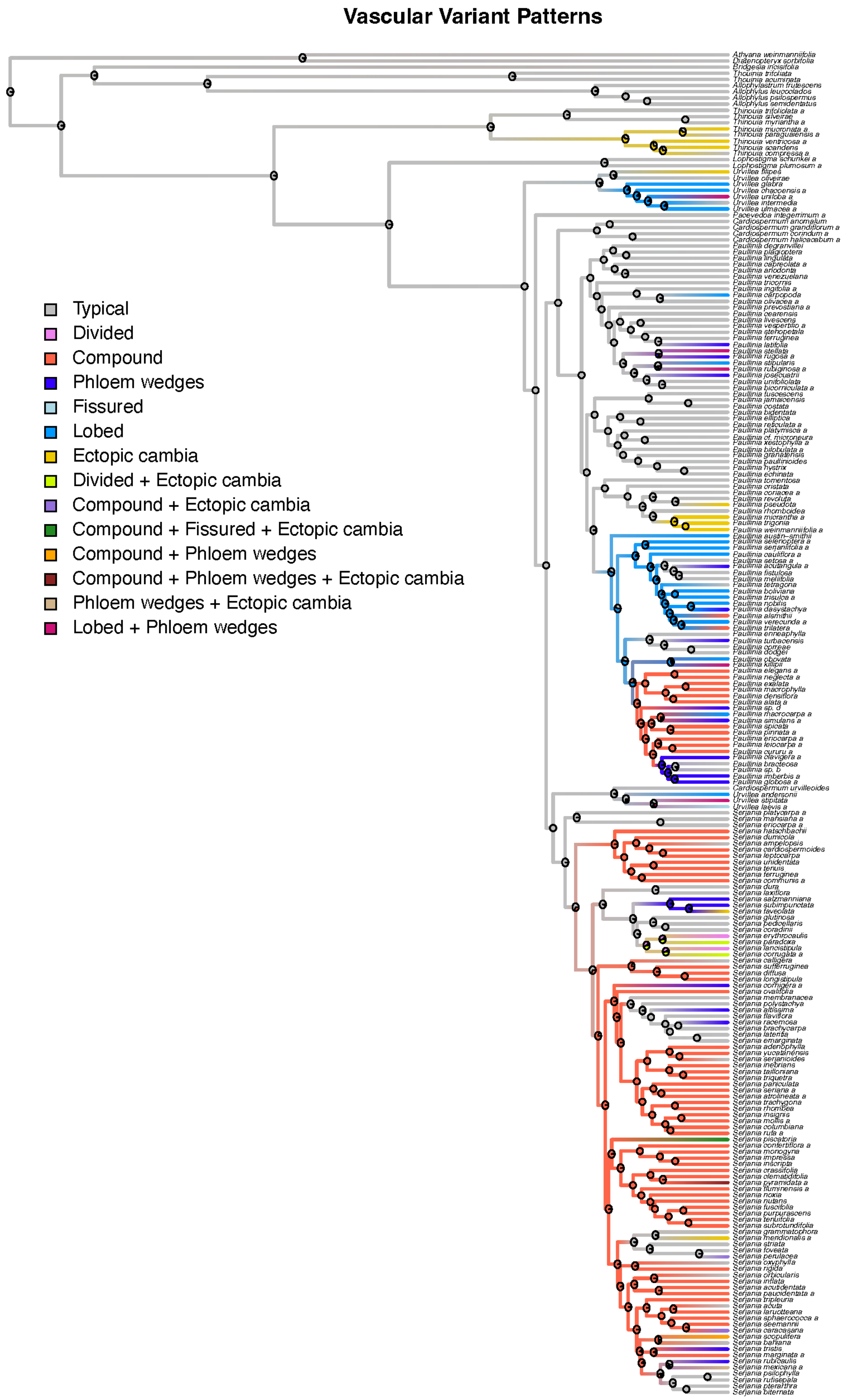


**Figure S11.** Stochastic character mapping illustrating the inferred evolutionary history of vascular variants across the three *categories* of vascular variants in 211 species of Paullinieae, along with nine outgroups. This result is based on 1000 simulated stochastic histories using the symmetrical (SYM) model. The ancestral node of all major lineages is reconstructed as a typical growth.

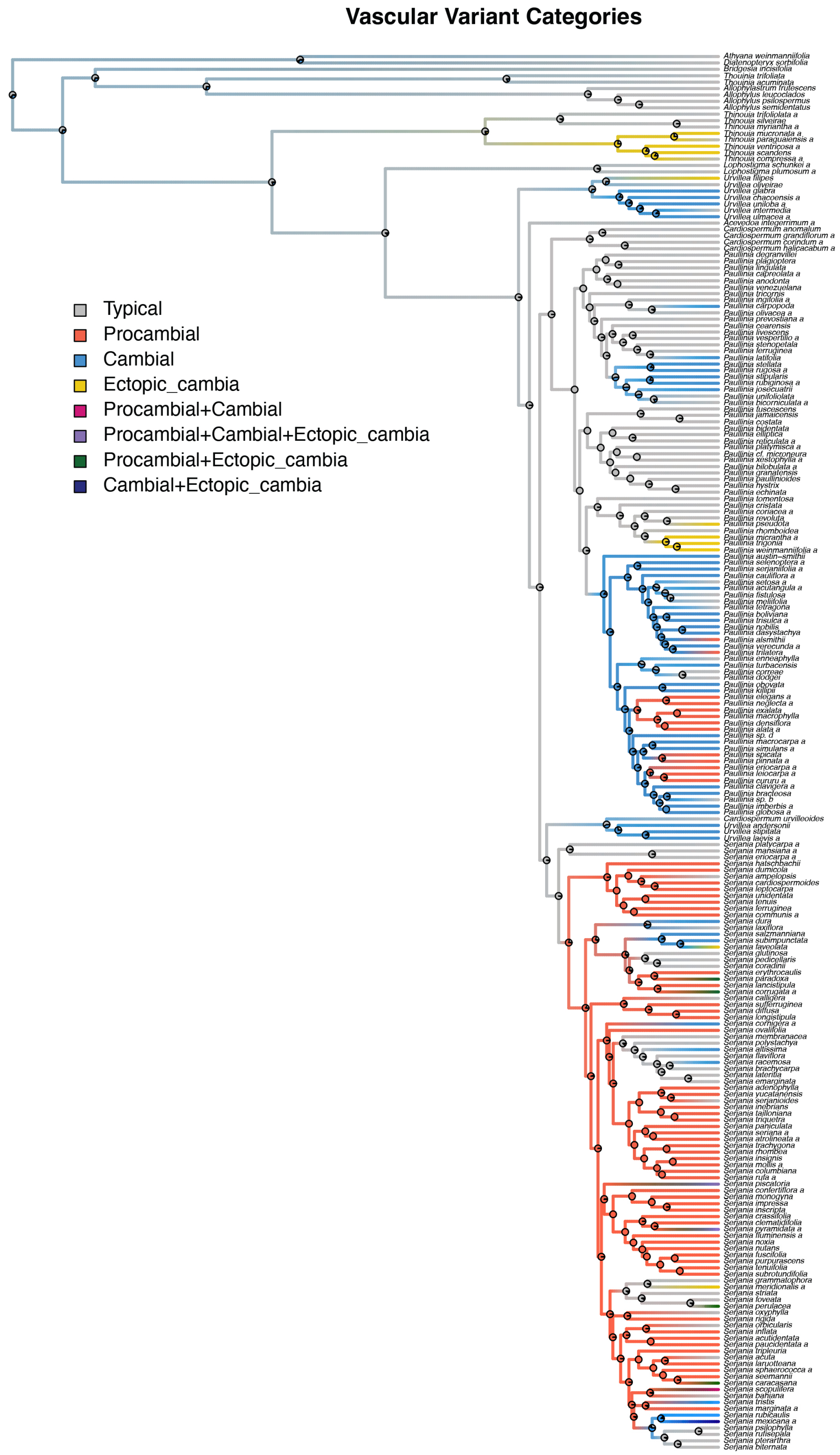


**Figure S12.** Stochastic character mapping illustrating the inferred evolutionary history of vascular variants across the six *patterns* of vascular variants in 211 species of Paullinieae, along with nine outgroups. This result is based on 1,000 simulated stochastic histories using the equal rates (ER) model. The ancestral node of all major lineages is reconstructed as a typical growth.

**Table S2**. Summary of target and final sampling for the Paullinieae phylogeny.

| Taxa* | # species | # species sampled | # samples | # species sampled | # silica specimens | # herbarium voucher specimens | # sequence samples | # total samples in tree |
|---|---|---|---|---|---|---|---|---|
| | Target sampling | | | Final sampling | | | | |
| *Pacevedoa*** | NA | NA | NA | 1 | 0 | 2 | 0 | 2 |
| *Cardiospermum* | 9? | 11 + 2 sp. | 24 | 5 | 1 | 11 | 0 | 12 |
| *Lophostigma* | 2 | 2 | 4 | 2 | 0 | 4 | 1 | 5 |
| *Paullinia* | 188 | 160 +10 sp. | 350 | 94 +2 sp. | 59 | 80 | 1 | 135 + 5 sp. |
| *Serjania* | 249 | 174 + 9 sp. | 353 | 100 +2 sp. | 37 | 89 | 1 | 125 + 2 sp. |
| *Thinouia* | 10 | 11+ 1 sp. | 24 | 8 | 0 | 16 | 1 | 17 |
| *Urvillea* | 18? | 12 +1 | 18 | 11 | 1 | 13 | 1 | 15 |
| *Outgroups* | - | 10 | 11 | 16 | 8 | 0 | 7 | 15 |
| **Total** | - | | **784** | **237 + 4 sp.** | **106** | **215** | **12** | **326+7 sp. = 333** |

*All genus names are *sensu lato*, including paraphyletic *Cardiospermum* and *Urvillea.*
**Taxa not recognized at the time of study design.

PNAS

**Table S5.** Number of species in each clade and estimated species count per state used in HiSSE analysis of Paullinieae and outgroups.

| Genera | # species per genus | # without vascular variants | # with vascular variants | # tree species | # liana species | # non-tendrilled species | # tendrilled species | # zygomorphic species | # actinomorphic species |
|---|---|---|---|---|---|---|---|---|---|
| **Paullinieae** | | | | | | | | | |
| *Pacevedoa* | 2 | 2 | 0 | 0 | 2 | 0 | 2 | 2 | 0 |
| *Cardiospermum* | 6 | 6 | 0 | 0 | 6 | 0 | 6 | 6 | 0 |
| *Lophostigma* | 2 | 2 | 0 | 0 | 2 | 0 | 2 | 2 | 0 |
| *Paullinia* | 179 | 118 | 61 | 0 | 179 | 0 | 179 | 179 | 0 |
| *Serjania* | 241 | 88 | 153 | 0 | 241 | 0 | 241 | 241 | 0 |
| *Thinouia* | 13 | 8 | 5 | 0 | 13 | 0 | 13 | 0 | 13 |
| *Urvillea* | 20 | 9 | 11 | 0 | 20 | 0 | 20 | 20 | 0 |
| **Subtotal** | **463** | **233** | **230** | **0** | **463** | **0** | **463** | **450** | **13** |
| **Outgroups** | | | | | | | | | |
| *Allophylastrum* | 1 | 1 | 0 | 1 | 0 | 1 | 0 | 0 | 1 |
| *Allophylus* | 212 | 212 | 0 | 212 | 0 | 212 | 0 | 212 | 0 |
| *Athyana* | 1 | 1 | 0 | 1 | 0 | 1 | 0 | 1 | 0 |
| *Bridgesia* | 1 | 1 | 0 | 1 | 0 | 1 | 0 | 1 | 0 |
| *Diatenopteryx* | 2 | 2 | 0 | 2 | 0 | 2 | 0 | 2 | 0 |
| *Thouinia* | 27 | 27 | 0 | 27 | 0 | 27 | 0 | 27 | 0 |
| **Subtotal** | **244** | **244** | **0** | **244** | **0** | **244** | **0** | **243** | **1** |
| **TOTAL** | **707** | **477** | **230** | **244** | **463** | **244** | **463** | **693** | **14** |

**Table S6.** Structure of HiSSE models used to test trait-dependent diversification.

| Model | Hidden States | Trait Dependent? | Notes |
|---|---|---|---|
| 1. Dull-null | 0 | No | Assumes no diversification rate variation or trait influence. Turnover and extinction fraction are equal across observed states ($\tau_1 = \tau_2$; $\varepsilon_1 = \varepsilon_2$). |
| 2. BiSSE | 0 | Yes | Trait-dependent diversification rates directly depend on the two states of the focal trait (without hidden states). Turnover differs between trait states ($\tau_1 \neq \tau_2$) but extinction fraction is equal ($\varepsilon_1 = \varepsilon_2$). |
| 3. HiSSE | 2 | Yes | Tests whether diversification is driven by observed trait and/or hidden factors. All turnover rates differ ($\tau_1$, $\tau_2$, $\tau_3$, $\tau_4$); extinction fraction is equal across all states ($\varepsilon_1 = \varepsilon_2 = \varepsilon_3 = \varepsilon_4$). |
| 4. CID-2 | 2 | No | Character-independent process in which diversification rates depend solely on the two states of an unobserved character (two hidden states; same number of parameters as BiSSE); allows diversification rate to vary across the phylogeny, but independently from the focal trait. Turnover varies across hidden states only ($\tau_1 = \tau_2$; $\tau_3 = \tau_4$); extinction fraction is equal ($\varepsilon_1 = \varepsilon_2 = \varepsilon_3 = \varepsilon_4$). |
| 5. CID-4 | 4 | No | Character-independent process in which diversification rates depend on the states of four hidden states (four hidden states; same number of parameters as HiSSE); allows diversification rate to vary across the phylogeny, but independently from the focal trait. Turnover varies across hidden states ($\tau_1 \neq \tau_2 \neq \tau_3 \neq \tau_4$); extinction fraction is equal ($\varepsilon_1 = \varepsilon_2 = \varepsilon_3 = \varepsilon_4$). |

**Table S7.** Summary of traits studied in the HiSSE analysis, with coding scheme, sampling fraction (i.e., proportion of species with each character state), and best model.

| Traits | Coding | Sampling fraction* | Best model |
|---|---|---|---|
| Tendrils | 0 = non-tendrilled<br>1 = tendrilled | 9/244<br>209/463 | HiSSE |
| Flower symmetry | 0 = actinomorphic<br>1= zygomorphic | 9/14<br>209/693 | CID-4 |
| Growth forms | 0 = tree or shrub<br>1 = liana | 9/244<br>209/463 | HiSSE |
| Vascular variants | 0 = typical growth<br>1 = vascular variants | 93/477<br>125/230 | CID-4 |

*Notation: species with the state in phylogeny/estimated number of species with the state in Paullinieae + outgroups. The estimated number of species within Paullinieae + outgroups is indicated in Table S5.

**Table S8.** Comparison of different HiSSE model fittings. The best-fitted model was selected based on the lowest AIC and the highest AICw (first model). The first row indicates the preferred model (lower AICc). Bold models with AICw > 0 (in bold) from the second series for flower symmetry and vascular variants were utilized for model averaging. loglik: the value of the maximum (negative) log-likelihood, AIC: Akaike information criterion, AICw: AIC weights, standardizing the AIC scores of fitted models.

| Growth forms | | | |
|---|---|---|---|
| | loglik | AICc | AICw |
| HiSSE.2 | -600.246580474043 | 1221.55596288045 | 1 |
| CID4.2 | -637.9193469167 | 1292.52768904871 | 0 |
| CID2.2 | -654.620260618715 | 1321.63862550283 | 0 |
| BiSSE_like.2 | -663.914357900746 | 1338.11173466942 | 0 |
| Null.2 | -681.667543934805 | 1371.52288129684 | 0 |

| Climbing mechanism | | | |
|---|---|---|---|
| | loglik | AICc | AICw |
| HiSSE.2 | -600.246580474043 | 1221.55596288045 | 1 |
| CID4.2 | -637.9193469167 | 1292.52768904871 | 0 |
| CID2.2 | -654.620260618715 | 1321.63862550283 | 0 |
| BiSSE_like.2 | -663.914357900746 | 1338.11173466942 | 0 |
| Null.2 | -681.667543934805 | 1371.52288129684 | 0 |

| Flower symmetry | | | |
|---|---|---|---|
| | loglik | AICc | AICw |
| **CID4.2** | **-642.340364157171** | **1301.36972352965** | **0.98** |
| **CID2.2** | **-648.447716705352** | **1309.29353767611** | **0.019** |
| **HiSSE.2** | **-646.418881705966** | **1313.9005653443** | **0.002** |
| BiSSE_like.2 | -660.198265292718 | 1330.67954945336 | 0 |
| Null.2 | -693.434160815898 | 1395.05611505903 | 0 |

| Vascular variants | | | |
|---|---|---|---|
| | loglik | AICc | AICw |
| **CID4.2** | **-754.564735885201** | **1525.81846698571** | **0.994** |
| **HiSSE.2** | **-757.512864102951** | **1536.08853013827** | **0.006** |
| CID2.2 | -767.655490281512 | 1547.70908482843 | 0 |
| BiSSE_like.2 | -781.178185450872 | 1572.63938976967 | 0 |
| Null.2 | -803.095044418298 | 1614.37788226383 | 0 |

**Table S9.** Collector and specimen information for each panel of figures in the manuscript. The table lists the collectors and corresponding specimen numbers for each figure panel.

| Collector Information for Images from Figures | | |
|---|---|---|
| **Figure Number** | **Panel** | **Collector Name & #** |
| Figure 1 | B | Acevedo-Rodríguez 15223 |
| | C | Acevedo-Rodríguez 16365 |
| | D | Acevedo-Rodríguez 17025 |
| | E | Acevedo-Rodríguez 17004 |
| | F-G | Acevedo-Rodríguez 16855 |
| | H | Acevedo-Rodríguez 3729 |
| | I-J | Acevedo-Rodríguez 16975 |
| | K | Acevedo-Rodríguez 1350 |
| | L-M | Somner 1794 |
| | N | Reproduced from Cunha Neto et al. (2023) |
| Figure 2 | B | Acevedo-Rodríguez 6568 |
| | C | Acevedo-Rodríguez 17136 |
| | D | Acevedo-Rodríguez 4641 |
| | E | Acevedo-Rodríguez 16873 |
| | F | Acevedo-Rodríguez 7650 |
| Figure 3 | C | Schunke 2310 |
| | D | Acevedo-Rodríguez 1734 |
| | E | Acevedo-Rodríguez 3719 |
| | F | Acevedo-Rodríguez 7535 |
| | G (top) | Acevedo-Rodríguez 3701 |
| | G (middle) | Acevedo-Rodríguez 1619 |
| | G (bottom) | Reproduced from Cunha Neto et al. (2018) |
| | H (top) | Acevedo-Rodríguez 6679 |
| | H (bottom) | Acevedo-Rodríguez 3296 |
| | I | Acevedo-Rodríguez 3713 |
| Figure S8 | A-D | Acevedo-Rodríguez 4655 |
| | E | Chery 23 |
| | F-G | Chery 45 |
| | H | Acevedo-Rodríguez s/n |
| | I-L | Acevedo-Rodríguez 3727 |

| | | |
|---|---|---|
| Figure S9 | A | Acevedo-Rodríguez 1540 |
| | B | Acevedo-Rodríguez 1561 |
| | C | Acevedo-Rodríguez 3691 |
| | D | Acevedo-Rodríguez 14420 |
| | E | Acevedo-Rodríguez 1573 |
| | F | Reproduced from Rizzieri et al. (2021) |
| | G | Acevedo-Rodríguez 4641 |
| | H | Acevedo-Rodríguez 14441 |
| | I | Acevedo-Rodríguez 15057 |
| | J | Acevedo-Rodríguez 5838 |
| | K | Guedes 31002 |
| | L | Acevedo-Rodríguez 16965 |
| | M | Reproduced from Marques et al. (2025) |
| | N | Acevedo-Rodríguez 16762 |
| | O | Neusa Tamaio s/n |
| Figure S10 | A | Acevedo-Rodríguez 16773 |
| | B | Reproduced from Rizzieri et al. (2021) |
| | C | Acevedo-Rodríguez s/n |
| | D | Acevedo-Rodríguez 14899 |
| | E | Acevedo-Rodríguez 15189 |
| | F | Acevedo-Rodríguez 15081 |
| | G | Marques 26 |
| | H | Marques 27 |
| | I-J | Acevedo-Rodríguez 3699 |

### Methods

#### 1. Sampling

**Leaf collection.** All collected samples met the following criteria: they were from collections made after 1990, consisted of green leaves, and were never collected in ethanol. Sampling was conducted in consultation with Dr. Pedro Acevedo-Rodríguez (curator of Sapindaceae). It represented collections by the top taxonomists in Paullinieae, namely Genise V. Somner, Maria S. Ferrucci, and Pedro Acevedo-Rodríguez, to ensure species verification.

**Species name curation.** To ensure that all names were spelled correctly, we used a species list curated by the author and specialist in the group, Dr. Acevedo-Rodríguez. We compared the spelling of the names with the Plants of the World Online (POWO) database (https://powo.science.kew.org/) (*SI Appendix,* Table S2). Along with accurate spelling, we utilized the POWO database as a gold standard for authority names and spelling, as it has been proven to be one of the best and most accurate databases for plant nomenclature (6).

#### 2. DNA analysis

**DNA Extraction and Quality Control.** Autogen can extract data from 384 specimens in a single run and has proven helpful in the 2019 publication on the *Paullinia* phylogeny (7). We conducted separate silica and herbarium voucher-batch runs to prevent contamination. DNA molecular weight was assessed using gel electrophoresis (3 µL of full-concentration DNA in a 1% agarose gel with a high molecular weight ladder). DNA 260/230 and 260/280 values were evaluated using a BioTek Epoch Microplate Spectrophotometer (Agilent) at the Smithsonian Laboratory of Analytical Biology. The total DNA was quantified via a spectrofluorimetric assay using a Qubit 4 Fluorometer (Life Technologies, Carlsbad, CA, USA). DNA quantification was assessed using a plate fluorometer SpectraMax M2 using the QuantiFluor reagent (Promega) at Cornell University Institute of Biotechnology. Further quality control and library construction were processed by Daicel accordingly: *Quality Control:* The gDNA was visually inspected and resuspended, and the total DNA was quantified via a spectrofluorimetric assay (Qubit 4). Samples were visualized using the TapeStation 4200 (Agilent) platform with a High Sensitivity D1000 tape to determine the molecular weight of each extraction.

**Library preparation.** Samples with high molecular weight DNA were handled with the Standard DNA library preparation protocol. Up to 80% of the gDNA was purified with a bead-based method and prepared into dual-indexed Illumina-compatible libraries, targeting an average insert size of approximately 500nt. Samples with low molecular weight DNA were handled with the Degraded DNA library preparation protocol. Up to 80% of the gDNA was taken through a blunt-end library preparation method that produces dual-indexed Illumina-compatible libraries. Samples with mixed molecular weight DNA were sonicated to produce inserts of approximately 500nt before following the Degraded DNA library preparation protocol. The indexed libraries were quantified with a spectrofluorimetric assay. Bait Capture: Capture pools were prepared from 6 to 12 libraries per reaction, pooled based on the input mass of gDNA during library preparation. Each capture pool was dried down to 7 µl by vacuum centrifugation. Captures were performed according to the myBaits v5.02 protocol, using myBaits Expert Angiosperms-353 baits with overnight hybridization and washes at 65°C (Standard) or 62°C (Degraded). Post-capture, the reactions were amplified and quantified again with a spectrofluorimetric assay. The captures were pooled in approximately equimolar ratios.

**Sequence assembly and filtering.** Following sequencing on the Illumina NovaSeq 6000 platform on a partial S4 PE150 lane, raw demultiplexed sequence reads were checked for quality using FastQC v0.11.9 (8). Adapters and low-quality bases were trimmed using Trimmomatic v0.39 with the following parameters: removal of adapter sequences using ILLUMINACLIP (2:30:10), trimming of leading and trailing bases below a quality score of 3, a sliding window of 4 bases cutting when the average quality dropped below 15, and discarding reads shorter than 36 bp. (9).

For enhanced locus recovery, we generated a custom target file with sequences for the 353 loci from 139 samples of Sapindaceae, downloaded from the Kew Tree of Life portal (https://treeoflife.kew.org). Hybpiper was executed with BWA (10), and the intronerate option enabled recovery of flanking introns and supercontigs, which include both exons and overlapping introns. The hybpiper_stats.py script was modified to generate a gene recovery report and a heatmap (02.hybpiper_stats.sh and 03.hybpiper_heatmap.sh). Paralogous and putatively chimeric sequences were identified and removed from subsequent analyses by concatenating the regions listed in the genes_with_long_parogs.txt and genes_derived_from putative_chimeric_stitched_contig.txt output files and removing all such problematic loci from each sample using the 04.hybpiper_paralog.sh script.

**Time-Calibrated Phylogeny.** One hundred randomly sampled bootstrap trees with the same topology but varying lengths were used as input for the treePL analysis. The following twelve outgroups were retained: *Allophylus semidentatus, A. psilospermus, A. leucoclados, Allophylastrum frutescens, Athyana weinmanniifolia, Blomia prisca, Bridgesia incisifolia, Diatenopteryx sorbifolia, Guindilia trinervis, Talisia nervosa, Thouinia acuminata,* and *Thouinia trifoliata. Cupania guatemalensis, C. racemosa,* and *Talisia cerasina* were excluded. For the treePL analysis, the best optimization parameters were determined using the "prime" command, and the smoothing parameter was determined through a cross-validation analysis with a cv start =100000 and cv stop =0.000000000001. Minimum and maximum dates were assigned to three nodes: (1) The root (*Talisia nervosa* + the entire tree) was set at 62–76 Mya based on secondary calibrations in (1); (2) *Thouinia* spp. + *Allophylus* spp. was set at 46–51 Mya based on a fossil-informed secondary calibration point from Joyce et al. (1). (3) *Urvillea* I, II, *Paullinia, Serjania, Cardiospermum* I, II, III was set at 18-19 Mya based on *Amperilorhiza heteroxylon* root fossil, which was phylogenetically placed through wood anatomy characters (2). Each tree was time-calibrated using the following parameters: numsites = 213409, nthreads = 4, and smooth = 0.000000001. Trees were visualized in FigTree (http://tree.bio.ed.ac.uk/software/figtree/).

#### 3. Plant micro technique

Specimens for developmental anatomy were obtained from two primary sources: liquid-preserved specimens collected in the field and dried plant collections (e.g., wood collections; USw) and herbaria (US, NY). Selection criteria for the specimens focused on obtaining stems at different developmental stages, specifically targeting young stems (ideally with shoot apical meristem) and mature stems with full secondary growth (including the presence of vascular variants when present). Specimens from herbaria were selected from species with more than three vouchers available in the collections, and stem samples were obtained through destructive sampling at the base of the stem in the voucher using a small saw. Samples were used to create temporary or permanent slides for anatomical investigation, following procedures adapted to the sample type, mostly based on original methods as summarized in the IAWA List of Microscopic Bark Features (11).

Initially, most samples were sectioned by hand using a razor blade to determine their developmental stage before further processing. Some young stems (with primary or early secondary growth near the shoot apex) were determined to be eligible for hand sectioning, given the integrity of the material and ease of sectioning. Particularly delicate samples were further embedded in Paraplast (Leica Biosystems) or Historesin (Leica Biosystems), then sectioned on a rotary microtome (Olympus CUT 4060). For adult stems (thick stems with secondary growth) from liquid-preserved specimens and samples from wood collections and herbaria, the preparation process involved boiling them in a 1:1 mixture of water and glycerin until the specimens could be easily cut with a disposable blade or submerged, respectively. This preparation step was crucial for ensuring the specimens were adequately rehydrated or softened for subsequent sectioning. For the larger specimens, following the boiling procedure, the samples were embedded in Polyethylene Glycol 1500 (PEG 1500) and sectioned with a sliding microtome (Spencer 880). The smaller stems, after the boiling procedure, were sectioned either by hand using a razor blade or through embedding and sectioning on a rotary microtome, as described above. Samples processed in Paraplast and PEG 1500 were stained with SafraBlau (a combination of Astra Blue and Safranin), whereas samples processed with historesin were stained with Toluidine Blue. Permanent slides were mounted with Eukitt. Slides were imaged using a light microscope (Olympus BH2) equipped with a digital camera (Amscope MU1000).

### 4. Character coding

**Terminology.** A recent review discussing the presence of vascular variants in Sapindaceae reported 10 possible stem or root architectures in Paullinieae lianas (12). In contrast, our study described only six patterns, as some patterns previously recognized (12) were merged with others, given that they follow a similar developmental pathway and were here treated as subtypes. That is the case, for instance, of “corded stem” and “neoformations” that were treated as subtypes of ectopic cambia (=successive cambia) (13). Similarly, we merged the two patterns of compound stems recognized by (12) into a single pattern with two subtypes. Another pattern, “Divided vascular cylinder with the delayed formation of a central cylinder,” was reduced to a combination in our study, as the divided stem organization occurs in most cases and during most developmental time, with the central cylinder, when present, resulting primarily in the formation of ectopic cambia (as in other instances with combination of a primary pattern with the late addition of vascular tissues through ectopic cambia). We also aggregated “axial elements separated in plates” (as recognized by (5)) with typical growth, given that we did not find species exhibiting "axial elements separated in plates” forming plates due to wide rays derived from the interfascicular cambium, which were evenly distributed around the stem circumference without other narrower rays, as described by most lineages that are traditionally characterized as having this pattern.

**PARAMO.** We modified the PARAMO pipeline (14) to accommodate vascular variants’ developmental and hierarchical complexity. Vascular variants are described with two levels: categories and patterns. The three categories represent the higher developmental level with changes to the origin and activity of vascular meristems: the procambium, the cambium, and the rise of an additional cambium. Within each category, multiple patterns share similar developmental modifications at the meristem level. In this context, we took our matrix, scoring the categories and patterns for all species (used for stochastic character mapping; see table below “A”), and amalgamated the different patterns as states of their respective categories (table below). The three categories + typical growth was treated as the character states for the paramo analysis (see table below “B”). Then, we performed stochastic character mapping by fitting hidden Markov models using the corHMM function in *corHMM* (15) and simulating stochastic histories with the makeSimmap function in *phytools* (16).

| **A – Stochastic Character Mapping** | |
|---|---|
| **Categories** | |
| **States** | **Code** |
| Typical | 1 |
| Procambial | 2 |
| Cambial | 3 |
| Ectopic_cambia | 4 |
| Procambial+Cambial | 5 |
| Procambial+Cambial+Ectopic_cambia | 6 |
| Procambial+Ectopic_cambia | 7 |
| Cambial+Ectopic_cambia | 8 |
| **Patterns** | |
| **States** | **Code** |
| Typical | 1 |
| Divided | 2 |
| Compound | 3 |
| Phloem-wedges | 4 |
| Fissured | 5 |

| | |
|---|---|
| Lobed | 6 |
| Ectopic-cambia | 7 |
| Divided+Ectopic-cambia | 8 |
| Compound+Ectopic-cambia | 9 |
| Compound+Fissured+Ectopic-cambia | 10 |
| Compound+Phloem-wedges | 11 |
| Compound+Phloem-wedges+Ectopic-cambia | 12 |
| Phloem-wedges+Ectopic-cambia | 13 |
| Lobed+Phloem-wedges | 14 |

**B – PARAMO**

| **Typical** | **Procambial variants** | **Cambial variants** | **Ectopic cambia** |
|---|---|---|---|
| 1= Not present | 1= Not present | 1= Not present | 1= Not present |
| 2 = Present | 2= Divided | 2 = Phloem_wedges | 2 = Present |
| | 3 = Compound | 3 = Fissured | |
| | | 4 = Lobed | |
| | | 5 = Lobed + Phloem wedges | |

### SI References


1. E. M. Joyce, *et al.*, Phylogenomic analyses of Sapindales support new family relationships, rapid Mid-Cretaceous Hothouse diversification, and heterogeneous histories of gene duplication. *Front. Plant Sci.* **14** (2023).

2. N. A. Jud, S. E. Allen, C. W. Nelson, C. L. Bastos, J. G. Chery, Climbing since the early Miocene: The fossil record of Paullinieae (Sapindaceae). *PLoS ONE* **16**, e0248369 (2021).

3. Y. C. Rizzieri, *et al.*, Ontogeny of divided vascular cylinders in *Serjania*: the rise of a novel vascular architecture in Sapindaceae. *IAWA J.* **42**, 121–133 (2021).

4. N. F. Marques, I. L. Cunha Neto, L. A. Brito, G. V. Somner, *Serjania piscatoria* (Paullinieae, Sapindaceae) as a symbol of vascular variants polymorphism. *Botanical Journal of the Linnean Society* **208**, 58–74 (2025).

5. P. Acevedo-Rodríguez, Lianas and climbing plants of the Neotropics. (2015). Available at: https://naturalhistory.si.edu/research/botany/research/lianas-and-climbing-plants-neotropics.

6. D. Costa, *et al.*, The big four of plant taxonomy – a comparison of global checklists of vascular plant names. *New Phytologist* **240**, 1687–1702 (2023).

7. J. G. Chery, P. Acevedo-Rodríguez, C. J. Rothfels, C. D. Specht, Phylogeny of *Paullinia* L. (Paullinieae: Sapindaceae), a diverse genus of lianas with dynamic fruit evolution. *Molecular Phylogenetics and Evolution* **140**, 106577 (2019).

8. S. Andrews, FastQC: A Quality Control Tool for High Throughput Sequence Data [Online]. (2010). Available at: http://www.bioinformatics.babraham.ac.uk/projects/fastqc/.

9. A. M. Bolger, M. Lohse, B. Usadel, Trimmomatic: a flexible trimmer for Illumina sequence data. *Bioinformatics* **30**, 2114–2120 (2014).

10. H. Li, R. Durbin, Fast and accurate short read alignment with Burrows-Wheeler transform. *Bioinformatics* **25**, 1754–1760 (2009).

11. V. Angyalossy, *et al.*, IAWA List of Microscopic Bark Features. *IAWA Journal* **37**, 517–615 (2016).

12. M. R. Pace, *et al.*, The wood anatomy of Sapindales: diversity and evolution of wood characters. *Braz. J. Bot* **45**, 283–340 (2022).

13. I. L. Cunha Neto, J. G. Onyenedum, Ectopic cambia: Connections between natural and experimental vascular mutants. *American J of Botany* **110**, e16246 (2023).

14. S. Tarasov, I. Mikó, M. J. Yoder, J. C. Uyeda, PARAMO: A Pipeline for Reconstructing Ancestral Anatomies Using Ontologies and Stochastic Mapping. *Insect Systematics and Diversity* **3**, 1 (2019).

15. J. M. Beaulieu, B. C. O'Meara, M. J. Donoghue, Identifying Hidden Rate Changes in the Evolution of a Binary Morphological Character: The Evolution of Plant Habit in Campanulid Angiosperms. *Systematic Biology* **62**, 725–737 (2013).

16. L. J. Revell, phytools: an R package for phylogenetic comparative biology (and other things). *Methods Ecol Evol* **3**, 217–223 (2012).
